# Aggregation pheromone 4-vinylanisole promotes the synchrony of sexual maturation in female locusts

**DOI:** 10.1101/2021.10.24.465628

**Authors:** Dafeng Chen, Li Hou, Jianing Wei, Siyuan Guo, Weichan Cui, Pengcheng Yang, Le Kang, Xianhui Wang

**Affiliations:** State Key Laboratory of Integrated Management of Pest Insects and Rodents, Institute of Zoology, Chinese Academy of Sciences, Beijing 100101, China; CAS Center for Excellence in Biotic Interactions, University of Chinese Academy of Sciences, Beijing 100049, China; Beijing Institutes of Life Sciences, Chinese Academy of Sciences, Beijing 100101, China

**Author notes:** Corresponding author: Dr. Xianhui Wang and Dr. Le Kang Institute of Zoology Chinese Academy of Sciences and. E-mail for other authors: Dafeng Chen Li Hou Siyuan Guo Weichan Cui Pengcheng Yang. These authors contribute equally to this paper (Dafeng Chen, Li Hou).

**Keywords:** sexual maturation, olfactory receptor, social interaction, endocrine hormone, pheromones

## Abstract

Reproductive synchrony generally occurs in various group-living animals; however, the underlying mechanisms remain largely unexplored. Here, we report that aggregation pheromone, 4-vinylanisole, plays a key role in promoting sexual maturation synchrony of female adults in the migratory locust, *Locusta migratoria*, a worldwide agricultural pest species that can form huge swarms with millions of individuals. Gregarious female locusts display significant synchrony of sexual maturation and oviposition whereas solitarious females and olfactory deficiency mutants do not. Only the presence of gregarious male adults can stimulate sexual maturation synchrony of female adults. Of the volatiles emitted abundantly by gregarious male adults, only 4-vinylanisole induces female sexual maturation synchrony, whereas this effect is abolished in mutants of 4- vinylanisole receptor. Interestingly, 4-vinylanisole mainly accelerates oocyte maturation of young females aged at middle developmental stages (3-4 days post adult eclosion). Juvenile hormone/vitellogenin pathway mediates female sexual maturation triggered by 4-vinylanisole. Our results highlight a “catch-up” strategy by which gregarious females synchronize their oocyte maturation and oviposition by time-dependent endocrinal response to 4-vinylanisole.

## Introduction

Reproductive synchrony, characterized by a pronounced temporal clustering of births, estrus, or mating, widely occurs in the animal kingdom, especially in group-living species (*Ims, 1990*). Several prominent cases are best known for their extreme manifestations, for example, sea turtle oviposition, firefly flashing, and fish spawning, involving a mass of individuals with the same reproductive state at certain time windows (*Buck and Buck, 1968, Harrison et al., 1984, Kelly and Sork, 2002*). Reproductive synchrony may offer adaptive advantages for group-living species, such as predation swamping and inbreeding avoidance (*Janzen, 1971*). Therefore, understanding how reproductive cycle is synchronized among individuals would provide insight to the biological flexibility in group-living animals.

Reproductive synchrony is a complex process that requires the integration of extra- and endo-signals to coordinate the timing of reproductive cycles between individuals in a group (*Kobayashi et al., 2002, Dey et al., 2015*). In fact, intra-group variation in developmental status can be induced by many factors, including different nutrition, temperature, and order of eclosion (*Ward and Webster, 2016*), which essentially makes synchronous reproduction between all members an apparent improbability. Social interaction is considerably critical for triggering reproductive synchrony of individuals in group-living species (*French and Stribley, 1985, Ims and Steen, 1990, Jovani and Grimm, 2008*). A well-known example is the Whitten effect which is induced by the presence of males in rodents, ewes, and monkey (*Vandenbergh, 1967, Cahill et al., 1974, Gattermann et al., 2002*). Various kinds of signals, odors, touch, or voice, can act as social clues to underpin synchronization with reproduction (*Rekwot et al., 2001, Kobayashi et al., 2002, Noguera and Velando, 2019*). Endo-signals, such as hormone release, gene expression and epigenetic modification, have also been suggested to be involved in these interaction processes (*Engel et al., 2016, Noguera and Velando, 2019*). However, the mechanisms by which social cue/hormone interaction synchronizes the reproductive cycles of individuals within local breeding groups remain largely unknown.

Locusts often form large swarms with hundreds to thousands of individuals, regarded as one of the most extraordinary examples of coordinated behavior (*Uvarov, 1977, Anstey et al., 2009*). Depending on population density, locusts display striking phenotypic plasticity, with a cryptic solitarious phase and an active gregarious phase (*Wang and Kang, 2014*). Gregarious locusts, compared to solitarious conspecifics, show much higher synchrony in physiological and behavioral events, such as egg hatching and sexual maturation, as well as synchronous feeding and marching behaviors (*Norris, 1954, Uvarov, 1977*). Some sort of vibratory stimulus, maternal microRNAs, and SNARE protein play important roles in the egg-hatching synchrony of gregarious locusts (*Chen et al., 2015, He et al., 2016, Nishide and Tanaka, 2016*). It has been revealed that the presence of mature male adults has effectively accelerating effects on synchrony of sexual maturation of immature male and female conspecifics in two locust species, *Schistocerca gregaria* and *Locusta migratoria* (*Norris, 1952, Loher, 1961, Guo and Xia, 1964, Norris, 1964*). Several kinds of mature-male-emitted volatiles have been proposed to accelerate sexual maturation of male adults in the desert locust (*Mahamat et al., 1993, Assad et al., 1997*). However, the pheromones that contribute to maturation synchrony of females have so far not been determined.

In this study, we investigated phase-related patterns and relevant mechanisms of sexual maturation of female adults in the migratory locust through multi-disciplinary studies, including physiology, chemical ecology, genomics, and gene manipulation. Unexpectedly, we found that aggregation pheromone, 4-vinylanisole, induces sexual maturation synchrony of female locusts. Our results highlight a parsimonious role of olfactory cues in the formation of locust swarms by triggering aggregation behavior and sexual maturation synchrony.

## Results

### Olfactory signals from gregarious male adults trigger the synchrony of female sexual maturation in locusts

We first investigated whether there was a difference of reproduction synchrony between gregarious and solitarious female locusts by determining the distribution of first oviposition date. The curve of the first oviposition time of gregarious females was much narrower than that of solitarious females (60% decrease in the standard deviation (SD), Figure 1A), implying that the first reproductive cycle was more consistent among gregarious female individuals. Then we measured the length of terminal oocyte relative to the final mature size, which is regarded as the indicator of female sexual maturation. The length of terminal oocyte increased with developmental stage in female adults of both phases. Gregarious female adults displayed more uniform and rapid patterns than that of solitarious females after 4 days post adult eclosion (PAE 4 days) (Figure 1B and Figure 1-figure supplement 1). These results indicate that gregarious female adults display significant sexual maturation synchrony and higher maturation rate.

**Figure 1.**
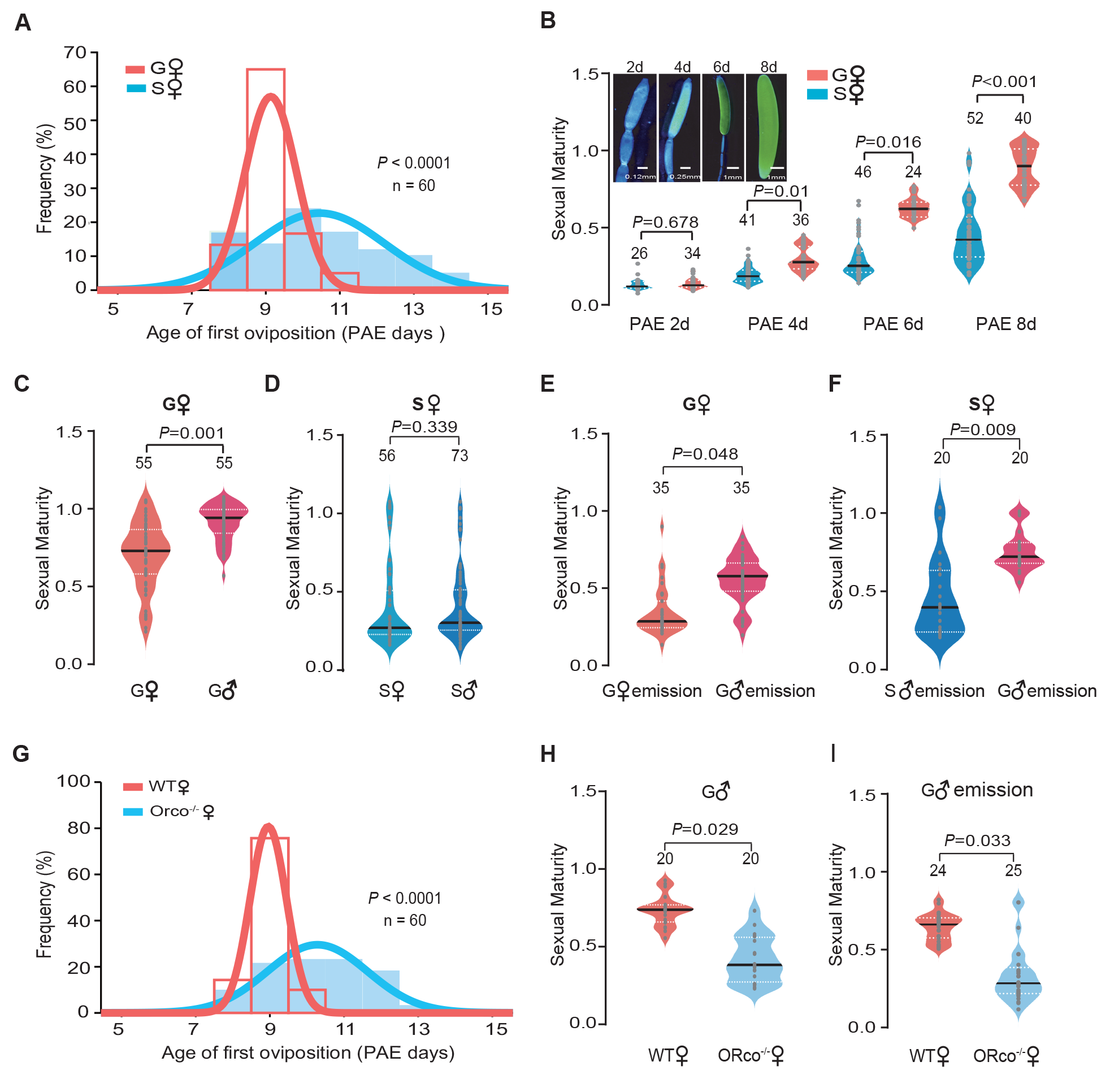
Olfactory signals from gregarious male adults trigger the synchrony of female sexual maturation in locusts. (A) Distribution of the first oviposition time of gregarious and solitarious phases. The first oviposition date was recorded after 6 days post adult eclosion (PAE 6 days) when individuals began to mate, and females that did not successfully mate with 24 h after pairing were excluded. Ages of first oviposition were indicated by days post eclosion. (B) The maturity of gregarious and solitarious females from PAE 2-8 days. The sexual maturity was presented as the length of terminal oocyte relative to the final mature size. (C) The maturity of gregarious females reared with gregarious males or females, separately. (D) The maturity of solitarious females reared with solitarious males or females, separately. (E) The maturity of gregarious females stimulated by volatiles released from gregarious males or females. (F) The maturity of solitarious females stimulated by volatiles released from gregarious or solitarious males. (G) Distribution of the first oviposition time in wild-type (WT) females and Orco female mutants (Orco^-/-^). (H) The maturity of WT females and Orco^-/-^ females reared with gregarious males. (J) The maturity of WT females and Orco^-/-^ females stimulated by volatiles released from gregarious males. Dark lines in violin plots indicate median value. White dotted lines indicate upper and lower quartile, respectively; Consistency analysis was analyzed using Levene’s test. The number of biological replicates and *P* values were shown in the figures.

We next investigated whether conspecific interactions can induce sexual maturation synchrony of female adults (Figure 1-figure supplement 2A and B). We found that the maturation synchrony of terminal oocytes of females was significantly retarded by the removal of male adults in gregarious phase (Figure 1C), but not in solitarious phase (Figure 1D). The exposure of odor blends from gregarious male adults significantly advanced the maturation synchrony of gregarious females and solitarious females, whereas no effects were observed when exposed to the background air, female odors, or odors from solitarious males (Figure 1E, F and Figure 1- figure supplement 2C, D).

To further explore the roles of olfactory cues in females’ sexual maturation process, we examined the performance of loss-of-function mutants of olfactory receptor co-receptor gene (Orco^-/-^) established by CRISPR/Cas9, which display significant olfactory deficiency (*Li et al., 2016*). The best-fit normal curve of the first oviposition date was much wider in Orco^-/-^ females than in wild-type females (WT), when they were reared together with gregarious males (with 63% increase in the SD, Figure 1G). When reared together with gregarious male adults or exposure to their odor blends, the sexual maturation of Orco^-/-^ females was less synchronous than that of WT females (Figure 1H, I and Figure 1-figure supplement 2E and F). Thus, olfactory signals from gregarious male adults are essential for triggering the synchrony of female sexual maturation in migratory locusts.

### 4-vinylanisole abundantly released by gregarious male adults mediates sexual maturation synchrony of female locusts

To identify key active compounds that can promote sexual maturation synchrony of female locusts, we determined the volatile emission dynamics of male adults in both phases from PAE 1- 8 days. In total, 14 chemicals were identified in the volatiles released by male adults (Figure 2A). Of them, 5 compounds displayed gregarious-male-abundant emission patterns, including phenylacetonitrile (PAN), guaicol, 4-vinylanisole (4-VA), vertrole, and anisole (Figure 2A and Figure 2-figure supplement 1), which might be potential active candidates.

**Figure 2.**
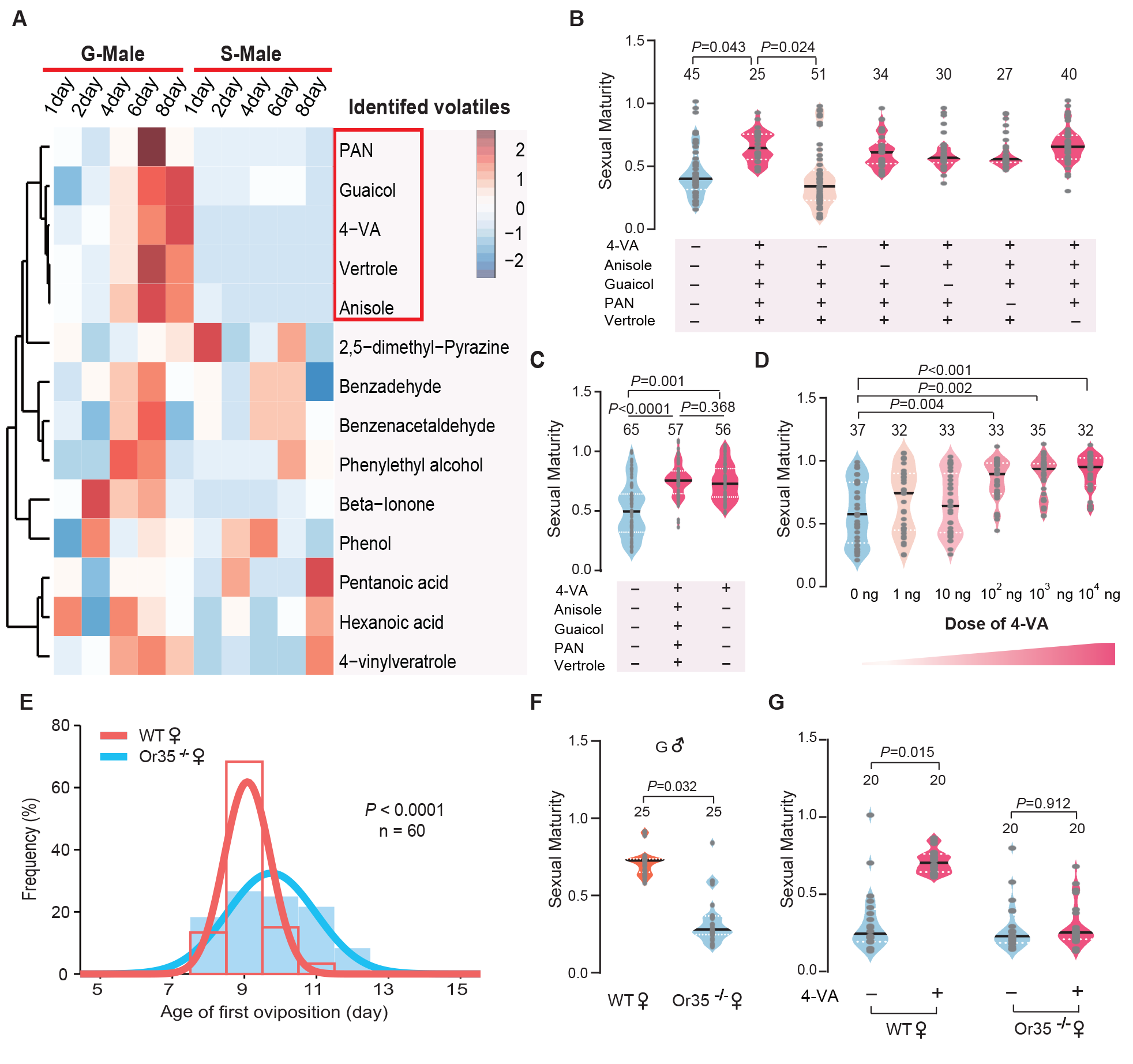
4-vinylanisole abundantly released by gregarious male adults promotes sexual maturation synchrony of female locusts. (A) Dynamic changes in volatiles released from male adults of two phases from PAE 1-8 days. (B) The maturity of gregarious females stimulated with different volatile mixtures containing gregarious male-abundant compounds. (C) The maturity of gregarious females treated with five kinds of gregarious male-abundant volatiles or 4-VA alone. (D) Dosage effects on the maturity of gregarious females after 4-VA stimulation. (E) Distribution of the first oviposition time in WT and Or35^-/-^ females. (F) The maturity of WT and Or35^-/-^ females reared with gregarious male adults. (G) The maturity of WT and Or35^-/-^ females with or without 4-VA stimulation. Dark lines in violin plot indicate median value. White dotted lines indicate upper and lower quartile, respectively; Consistency analysis was analyzed using Levene’s test. The number of biological replicates and *P* values were shown in the figures.

We exposed gregarious young female locusts for 6 days after fledging to different synthetic blends of those five compounds (PAN, guaicol, 4-VA, vertrole, and anisole). The full blend of five components was effective in promoting the synchrony of oocyte development. Only the omission of 4-VA, but not other four compounds, from the full blend lost the accelerating effects on sexual maturation synchrony of gregarious females (Figure 2B). Moreover, the exposure to 4- VA can induce similar effects on female sexual maturation synchrony to the full blend (Figure 2C). In addition, the accelerating effects of 4-VA on maturation synchrony displayed a dose-threshold pattern, with an effective concentration more than 100 ng (Figure 2D and Figure 2- figure supplement 2). We further examined the performance of Or35^-/-^ females that cannot sense 4-VA (*Guo et al., 2020*). Compared to WT females, the best-fit normal curve of the first oviposition date of Or35^-/-^ females was much wider (52% increase in the SD, Figure 2E). The sexual maturation in Or35^-/-^ females was more uneven than that of WT females when they were reared together with gregarious males (Figure 2F) or exposed to the odors of gregarious males (Figure 2-figure supplement 3). Moreover, the synchronous effects of 4-VA completely disappeared in Or35^-/-^ females (Figure 2G).

### 4-VA action needs a critical time window on sexual maturation synchrony in young females

In fact, intragroup variation generally exists in the maturation period of female locusts due to differences in nymph experience, nutrition, and fledging time (*Uvarov, 1977*). We hypothesized that there is a differential effect of 4-VA on the maturation rate for female individuals with distinct developmental statuses to achieve maturation synchrony. To test this, we determined the accelerating effects of 4-VA on young females at three different ages after fledging: PAE 1-2 days, 3-4 days, and 5-6 days, respectively. We found that female maturation synchrony was significantly enhanced only when young females were treated by 4-VA at PAE 3-4 days, while did not change at PAE 1-2 days or PAE 5-6 days, indicating the time window of 4-VA action on sexual maturation of females (Figure 3A). Moreover, we compared the effects of gregarious males with different ages on female maturation. The maturation synchrony of females was significantly enhanced by gregarious males aged at PAE 3-4 days and PAE 5-6 days (Figure 3- figure supplement 1), which could release more 4-VA (Figure 2-figure supplement 1). By contrast, rearing together with the fifth instar of gregarious males and male adults aged at PAE1-2 days did not significantly affect the maturation synchrony of gregarious female adults (Figure 3- figure supplement 1).

**Figure 3.**
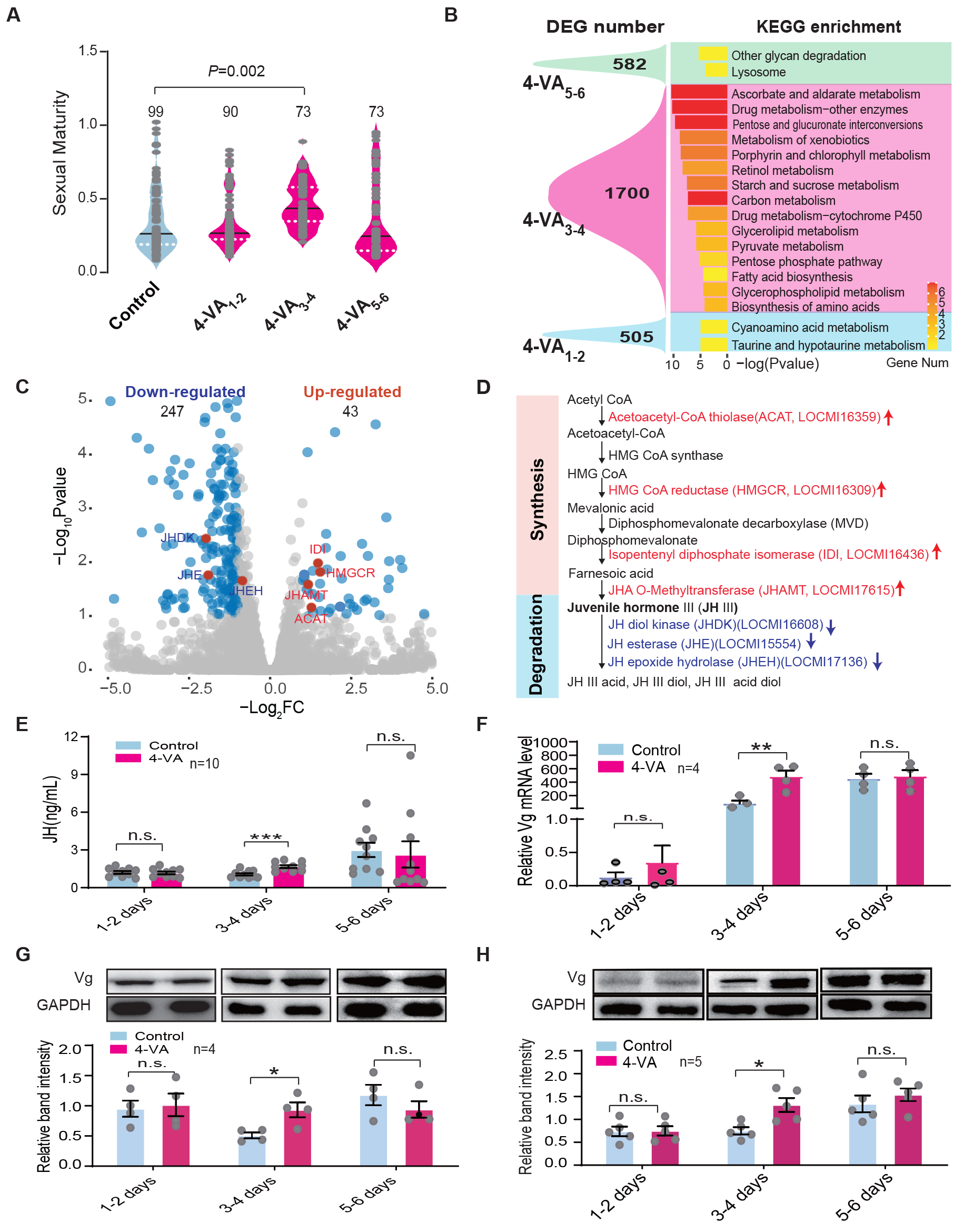
4-VA promotes sexual maturation synchrony in young females by enhancing JH/Vg signaling pathway at PAE 3-4 days. (A) The 4-VA effects on sexual maturity of gregarious females at different developmental stages. (B) Enrichment of DEGs and KEGG in the fat body of gregarious females after 4-VA stimulation at different developmental stages. (C) Volcano plot of RNA-seq in the CC-CA complex of gregarious females after 4-VA stimulation at PAE 3-4 days. Red dots indicate genes related to JH metabolism. (D) Expression changes of JH metabolism related genes in the CC-CA by 4-VA stimulation. Red and blue indicate upregulated and downregulated, respectively. (E) JH titers in the hemolymph, (F) The mRNA levels, (G) The protein levels of Vg in the fat body, and (H) the protein levels of Vg in the ovary of gregarious females after 4-VA stimulation at different developmental stages. Dark lines in violin plot indicate median value. White dotted lines indicate upper and lower quartile, respectively; columns show means ± SEM. Consistency analysis of maturity was analyzed using Levene’s test. The mRNA and protein levels were analyzed using Student’s t-test. The number of biological replicates and *P* values were shown in the figures. n.s. not significant.

Gene expression profiles in fat body tissue have been demonstrated to correlate tightly with the sexual maturation of female locusts (*Guo et al., 2014*). Therefore, we further evaluated the time-window effects of 4-VA on female sexual maturation at the transcriptomic level. Through RNA-seq, we verified that the gene expression profiles of fat body displayed more remarkable changes in female adults exposed to 4-VA at PAE 3-4 days (1700 differentially expressed genes (DEGs)) than those at PAE 1-2 days (505 DEGs) and at PAE 5-6 days (582 DEGs) (Figure 3B). Meanwhile, Kyoto encyclopedia of genes and genomes (KEGG) enrichment analysis showed that there were more signal pathways affected by 4-VA treatment at PAE 3-4 days than PAE 1-2 days and PAE 5-6 days. Notably, genes related to energy metabolism, such as retinol metabolism, glycerolipid metabolism, pyruvate metabolism, as well as fatty acid biosynthesis, which play essential roles in ovary development, were significantly activated by 4-VA treatment at PAE 3-4 days (Figure 3B). Thus, PAE 3-4 days should be a critical time window for 4-VA-induced acceleration of female sexual maturation.

### JH/Vg signaling pathway mediates the accelerating effect of 4-VA on sexual maturation synchrony in young females

To explore the regulatory mechanism underlying the time-window effects of 4-VA on the sexual maturation synchrony of female locusts, we examined the performance of major signaling pathways involved in the sexual maturation of female locusts. Firstly, we determined whether females display time-dependent electrophysiological response to 4-VA by performing EAG and SSR experiments. We found that 4-VA-induced EAG and SSR responses of female adults displayed obvious dose-dependent effects (Figure 3-figure supplement 2A-C). However, there was no difference of EAG and SSR responses of females among the ages of PAE 2 days, PAE 4 days, and PAE 6 days, although LmigOr35 expression levels were dynamic during ovary development (Figure 3-figure supplement 2D and 3). These results suggested that peripheral olfactory perception may not be involved in the time-window effects of 4-VA. We then compared gene expression profiles of two main neuroendocrinal tissues, the brain (Br) and corpus cardiacum-corpora allatum complex (CC-CA), between the controls and 4-VA-exposed females at PAE 3-4 days. Notably, gene expression profiles in CC-CA significantly changed upon 4-VA treatment, with 290 differentially expressed genes, much more than that in the brain (89 DEGs) (Figure 3C and Figure 3-figure supplement 4), implying the molecular and physiological activities in CC-CA might be remarkably affected by 4-VA stimuli. Moreover, a series of DEGs of CC-CA involved in juvenile hormone (JH) metabolism were enriched. There was significantly higher expression of genes related to JH synthesis but lower expression of genes associated with JH degradation (Figure 3C, D and Figure 3-figure supplement 5 and Supplementary file 1). These results indicated a potential role of JH signaling pathway in mediating the effects of 4-VA at PAE 3-4 days.

We therefore tested whether 4-VA exposure can affect the hemolymph JH titer in immature females. As expected, the JH titer was significantly elicited by 4-VA exposure in females aged at PAE 3-4 days, rather than PAE 1-2 days or PAE 5-6 days (Figure 3E). Similarly, the expression levels of vitellogenin (Vg), a key downstream component of JH signaling triggering ovary development in locusts (*Song et al., 2014*), were prominently increased in fat body and ovary of females aged at PAE 3-4 days upon 4-VA stimuli (Figure 3F-H). By comparison, JH titer did not significantly change in Or35^-/-^ females exposed to 4-VA, contrast to over two-fold increase in WT females (Figure 4A). Similar patterns were observed for the expression levels of Vg in fat body (Figure 4B and Figure 4-figure supplement 1A) and ovary (Figure 4C). To verify the critical roles of the JH/Vg pathway in mediating the effect of 4-VA, we further carried out rescue experiments by the injection of JH analog (methoprene) in Or35^-/-^ females. Methoprene-injected Or35^-/-^ females displayed more uniform sexual maturation (Figure 4D). Meanwhile, the expression levels of Vg in fat body and ovary, significantly increased in methoprene-injected Or35^-/-^ females (Figure. 4E, 4F, and Figure 4-figure supplement 1B). These results provide clear evidence that the JH/Vg signaling pathway can mediate the time-dependent accelerating effects of 4-VA on sexual maturation synchrony in female locusts.

**Figure 4.**
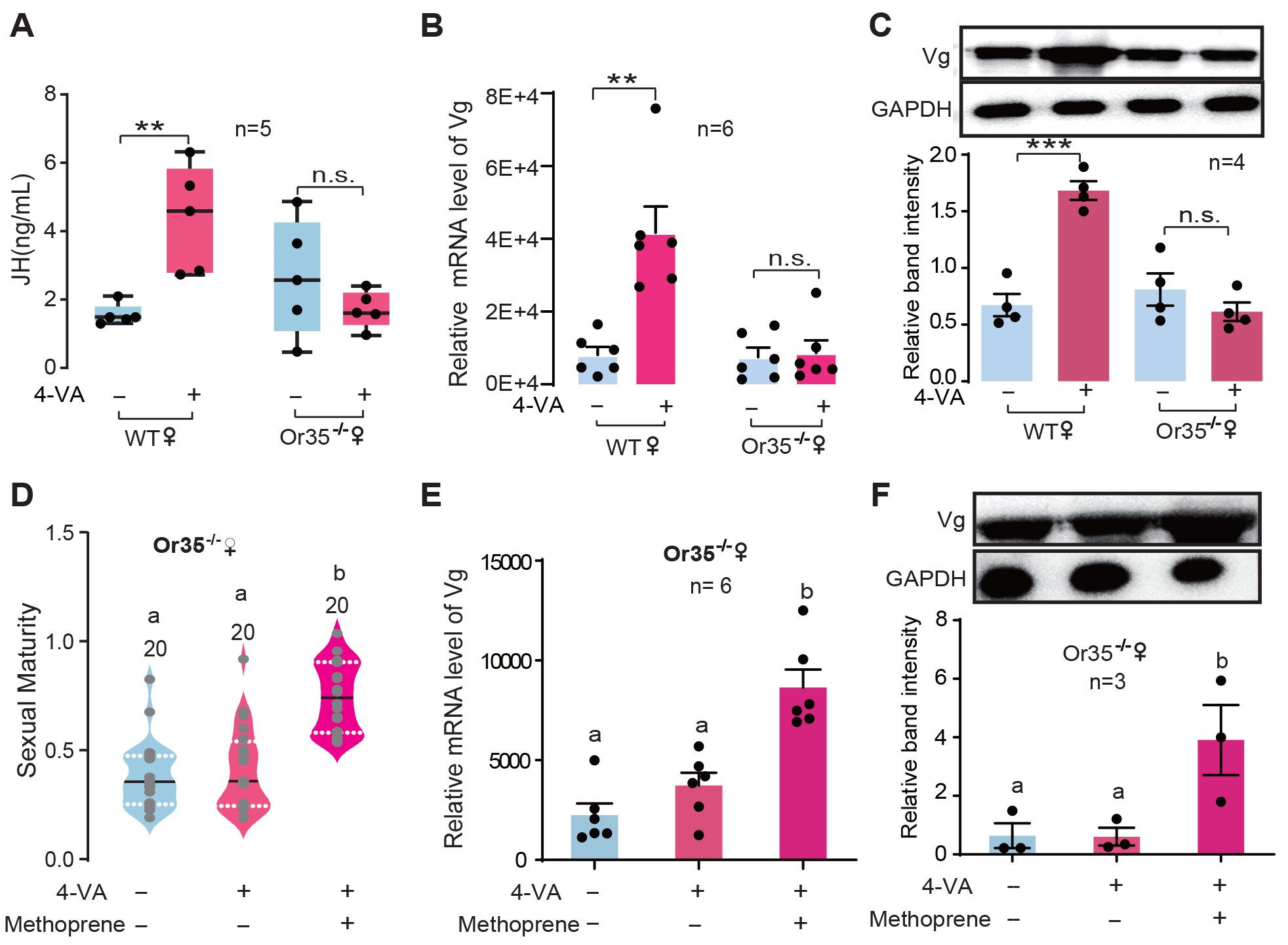
JH/Vg pathway indeed mediates the stimulatory effects of 4-VA on female sexual maturation. (A) JH titers in the hemolymph of WT and Or35^-/-^ females after stimulation by 4- VA at PAE 3-4 days. (B) The mRNA levels of *Vg* in the fat body and (C) protein levels of Vg in the ovaries of WT and Or35^-/-^ females after stimulation by 4-VA at PAE 3-4 days. (D) The effects of JH analog treatments on the maturity of Or35^-/-^ females exposed to 4-VA. (E) The mRNA level of *Vg* in the fat body and (F) protein level of Vg in the ovary of Or35^-/-^ females with 4-VA stimulation and JH analog treatments at PAE 3-4 days. Boxplots depict median and upper and lower quartile; Lines in droplet diagram indicate median value; columns show means ± SEM. One-way ANOVA, *P* < 0.05, Columns labeled with different letters indicate a significant difference between these groups. The number of biological replicates.

## Discussion

Our current study demonstrates conclusively that aggregation pheromone, 4-VA, acts to promote female maturation synchrony in locusts. The pheromone is abundantly released from gregarious male adults and speeds up oocyte development of females aged at PAE 3-4 days through activating JH synthesis and vitellogenesis (Figure 5). Our findings highlight a “catch-up” strategy of reproductive synchrony by a time-window effect combined with extra- and endo- signals in group-living animals.

**Figure 5.**
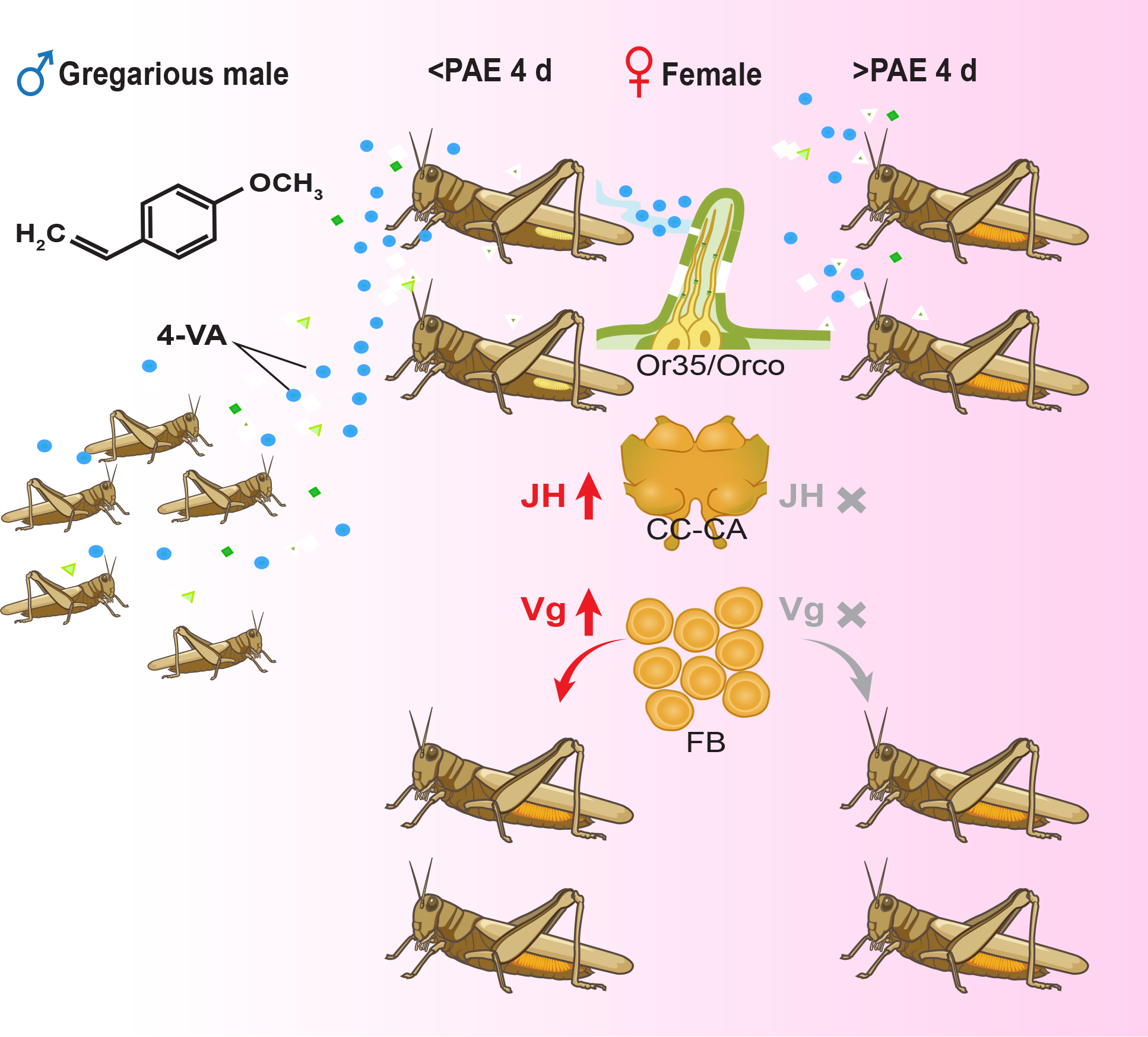
Schematic mechanisms underlying 4-VA-induced synchrony of female sexual maturation. 4-VA released from gregarious male locusts can significantly accelerate the ovary development of females with less-developed ovaries (approximately before PAE 4 days) but not well-developed ovaries (after PAE 4 days). Mechanistically, after recognition by Or35 expressed in antennae, 4-VA promoted JH synthesis in the CC and vitellogenesis in the fat body, thus accelerating female sexual maturation. The time-dependent stimulatory effects of 4-VA on ovary development finally lead to the synchrony of female sexual maturation. CC-CA, corpora cardiaca and corpora allata; FB, fat body.

We prefer that 4-VA acts as a critical multifunctional pheromone for the formation of large locust swarms. Earlier, we have demonstrated that 4-VA is mainly released by gregarious nymphs and male adults (*Wei et al., 2017*) and can induce strong attraction behavior of both gregarious and solitarious phases (*Guo et al., 2020*), indicating its releaser pheromone role in in keeping locust individuals living together. Meanwhile, the present study shows that 4-VA, acting as a primer pheromone, promotes the maturation synchrony of young female adults, which might facilitate simultaneous oviposition and egg hatching to reduce the predation risk of an individual via the dilution effect (*Ward and Webster, 2016*). Thus, a dual role of 4-VA, including both primer and releaser pheromones, could be proposed in triggering the formation of locust swarms. The maintenance and coordination of locust swarming require elaborate communication mechanisms behind the interaction among individuals (*Pener and Simpson, 2009, Wang and Kang, 2014*). It is likely an effective and optimized strategy of group-living animals to use a single chemical pheromone to elicit both behavioral and endocrine responses in conspecifics (*Rekwot et al., 2001*). The action of 4-VA displays a remarkable context-dependent manner, such as phase-, sex-, dose- and time-dependent, reflecting physiological adaption of locusts to the highly dynamic nature of population density.

We demonstrate clearly that crowding with older gregarious male adults could accelerate sexual maturation of female adults in the migratory locust, which is consistent with most of the previous finding in locusts (*Guo and Xia, 1964, Norris and Richards, 1964, Torto et al., 1994, Mahamat et al., 2000*). In fact, volatile pheromones released by mature males have for a long time been speculated to accelerate maturation of young adults in locusts (*Guo and Xia, 1964, Norris and Richards, 1964*). Several compounds of the volatiles from mature males have been proposed to accelerate the maturation of male adults in the desert locust (*Torto et al., 1994, Mahamat et al., 2000*). However, any distinct active accelerating compounds for sexual maturation of female adults in locusts until now. Here we show that a single minor component (4- VA) of the volatiles abundantly released by gregarious male adults is sufficient to induce the maturation synchrony of female adults. The possible explanation is that there might exit a sex- differential pattern of maturation-accelerating pheromones due to different selective pressures between two sexes in response to social environments. Different encoding strategies might have evolved: multicomponent pheromones for males and single active component for females. Further exploration will be performed to confirm this hypothesis by determining whether 4-VA has maturation-accelerating effects on male adults in the migratory locust in future.

A dose-dependent manner was found for the maturation synchrony effect of 4-VA. We find that only gregarious males aged after PAE 3 days have the accelerating effects on female maturation synchrony, which may be attributed to their significantly increased 4-VA content during adult development. Although gregarious nymphs (the fifth instar) and female adults can release relatively small amount of 4-VA (*Wei et al., 2017*), they did not promote female maturation based on our current results. Thus, the accelerating effects on female maturation synchrony induced by gregarious male adults may depend largely on 4-VA content they released. The ineffectiveness of the fifth nymphs and females in maturation acceleration of female adults may due to their low 4-VA content under efficient threshold. In fact, the fifth nymphs have been shown to display inhibiting effects on male maturation in *S. gregaria* (*Assad et al., 1997*). Therefore, the mechanisms underlying pheromone-mediated sexual maturation may differ between different locust species. Given that 4-VA has not been detected in *S. gregaria* (*Torto et al., 1996*), whether this volatile has maturation-accelerating effect in this locust species needs further validation.

Our results reveal that JH signaling pathway presents as the critical endocrinal factor mediating the accelerating effect of 4-VA on female maturation. This finding is consistent with the role of JH as the major gonadotropin modulating Vg biosynthesis in the fat body and its uptake by the growing oocytes in the migratory locust (*Jindra et al., 2013, Guo et al., 2014, Song et al., 2014*). It is also supported by the significance of CA (a major JH-biosynthesis tissue) in pheromone-induced maturation process in the desert locust (*Odhiambo, 1966*). Interesting, it has been suggested that the release of the maturation accelerating pheromone by adult males is under the control of CA (*Loher, 1960*). Thus, there should be a complex feedback interaction between 4-VA and JH signaling pathway. Extensive studies have established the central roles of JH signaling in mediating the effects of social interactions on reproduction in different kinds of insect species, including eusocial insects (*Robinson and Vargo, 1997, Korb, 2015*), the burying beetle, *Nicrophorus vespilloides* (*Engel et al., 2016*), the German cockroach, *Blattella germanica* (*Uzsak and Schal, 2012*) and so on. Such an interaction between social clues and internal hormonal signals that coordinates ovary development is also common among group-living vertebrates (*Drickamer, 1977, McClintock, 1978, Berger, 1992*).

We demonstrate that 4-VA stimulates sexual maturation of young females within a distinct developmental time window. Compared to the females aged at PAE 1-2 days and 5-6 days, the females aged at PAE 3-4 days were more sensitive to 4-VA stimuli. This point was strongly supported by several lines of evidence from temporal-dependent comparisons of oocyte development, gene expression profiles, JH titer, as well as Vg biosynthesis. It has been shown that JH titers, Vg expression, the size of terminal oocytes, dramatically increased at PAE 3-4 days, implies the PAE 3-4 days is an essential time window for JH-regulated ovary development in female locusts (*Luo et al., 2017, Wu et al., 2018*). The finding that 4-VA accelerates maturation of less developed females rather than more developed females supports a ‘catch-up’ model in achievement of female maturation synchrony in locusts. Peripheral and central neural sensitivity to olfactory clues have been demonstrated to vary with developmental stages or physiological statuses (*Guo et al., 2011, Gadenne et al., 2016*). Given this, sensory processing sensitivity or JH biosynthesis activity might be involved in the stage-dependent sensitivity of females to 4-VA stimuli. However, peripheral olfactory neuron might not be involved in the stage-specific sensitivity to 4-VA stimuli, because we did not detect significant changes of peripheral electrophysiological response during female ovary development. A possible explanation is that signaling factors responsible for JH synthesis might be turned on specifically at Mid-PAE of female locusts upon 4-VA stimuli, such as GPCRs and transcription factors (*Bendena et al., 2020*). Although there are only a few DEGs in the brain of females exposed to 4-VA, we cannot exclude the involvement of the central nerve system pathway by other regulatory mechanisms, for example, neurotransmitter release, or post-transcription regulation (*Nouzova et al., 2018*). Further studies should elucidate detailed mechanisms of the linking between 4-VA and JH biosynthesis in female locusts.

In summary, we revealed a catch-up strategy of female reproductive synchrony in locust swarms, whereby 4-VA acts as a maturation-accelerating pheromone hastening less developed females through triggering JH biosynthesis. Our findings provide novel insight into the mechanisms underlying individual interaction during aggregation in group-living animals.

## Materials and Methods

### Experimental insects

All insects used in experiments were reared in the same locust colonies at the Institute of Zoology, Chinese Academy of Sciences, Beijing, China. Briefly, gregarious locusts were reared in cages (30 cm × 30 cm × 30 cm) with 800 to 1000 first-instar insects per cage in a well-ventilated room. Solitarious locusts were individually raised in a ventilated cage (10 cm × 10 cm × 25 cm). All locusts were cultured under the following conditions: a L14:D10 photoperiod, temperature of 30 ± 2°C, relative humidity of 60 ± 5%, and a diet of fresh greenhouse-grown seedlings and bran.

### Oocyte length measurement

The ovary was dissected and placed in locust saline, and the terminal oocytes were isolated. The lengths of terminal oocytes were photographed and measured under the Leica DFC490 stereomicroscope (Leica, Germany). The sexual maturity was presented as the length of terminal oocyte relative to the final mature size.

### Recording of the distribution of first oviposition time in female locusts

Individuals between 0 and 24 hours after adult molting are referred as PAE 1 day adults, with each subsequent day representing an additional 24-hour period. Given that both gregarious and solitarious locusts begin to mate at PAE 6 days, the first oviposition time was recorded after PAE 6 days. For gregarious locusts, ten females and ten males at PAE 6 days were placed in a cage (30 cm × 30 cm × 30 cm). The females were individually marked, and their first oviposition times were recorded by collecting egg pods every 4 hours per day after mating. For solitarious locusts, each female was reared together with a single male at PAE 6 days, and the first oviposition time was recorded by collecting egg pods every day after mating. To avoid the effects caused by asynchronous mating, females that did not successfully mate within 24 h after paired rearing were excluded in both phases. The distribution curve of the first oviposition time was calculated based on data collected from all females.

### The effects of conspecifics interaction on female sexual maturation

Given that the difference of female sexual maturation synchrony between gregarious and solitarious phases appeared at PAE 6 days, the length of terminal oocytes was detected at PAE 6 days after each treatment in subsequent experiments. For the stimulation of gregarious females, ten gregarious females were reared with ten gregarious males or ten gregarious females in a same cage (15 cm × 15 cm × 10 cm) from PAE 1 days to 6 days (Fig S2A). For the stimulation of solitarious females, one solitarious females was reared with one solitarious male or one solitarious female in a same cage (15 cm × 15 cm × 10 cm) from PAE 1 days to 6 days (Fig S2B). The ovaries of treated females were dissected in locust saline and the lengths of terminal oocytes were measured as described above.

### The effect of locust volatiles on female sexual maturation

To determine the effect of locust volatiles on female sexual maturation, treated female adults were separately reared with females or male adults by a breathable partition. For gregarious phase, ten gregarious females were reared with ten gregarious males or ten gregarious females in a breathable partition cage (15 cm × 15 cm × 10 cm) from PAE 1 days to 6 days (Fig. S2C). For solitarious phase, one solitarious female was reared with ten gregarious males or ten solitarious males in a breathable partition cage (15 cm × 15 cm × 10 cm) from PAE 1 days to 6 days (Fig. S2D). The ovaries of treated females were dissected in locust saline and the lengths of terminal oocytes were measured as described above.

### Static solid phase microextraction (SPME) and GC-MS-MS

The volatiles of males and females in both phases at PAE 1, 2, 4, 6, 8 days were collected by SPME for 30 min following our previously study (*Wei et al., 2017*). In detail, a fiber (PDMS/DVB 65 μm) was introduced into a glass jar (10.5 cm high × 8.5 cm internal diameter) to absorb odors. The SPME volatiles collected from an empty glass jar for 30 min served as the control. Eight biological replicates were performed for each treatment. The fibers with absorbed odors were subjected to chemical analyses with GC-MS/MS. A Bruker GC system (456-GC) coupled with a triple quadrupole (TQ) mass spectrometer (Scion TQ MS/MS, Inc., German) equipped with an DB-1MS column (30 m × 0.25 mm ID × 0.25 μm film thickness, Agilent Technologies) was used to quantify the volatile compounds in the SPME samples. Bruker chemical analysis MS workstation (MS Data Review, Data Process, version 8.0) was used to analyze and process the data. Mixed samples consisting of standard compounds in different dosages (0.1, 1, 5, 10, and 20 ng/μl) were used as external standards to develop the standard curves to quantify the volatiles. The same thermal program and MRM method were used for standard compound detection.

### Odor treatment assay

For mixture treatment, ten gregarious females were stimulated by the mixed odor blend (the concentrations of PAN, Guaiacol, 4-VA, vertrole and Anisole were 1 μg/μl, 10 μg/μl μg/μl, 2 μg/μl, 3μg/μl, respectively) or paraffin oil from PAE 1 days to 6 days. In detail, a breathable vial containing the mixture or paraffin oil was placed with ten females in a cage for 6 days. The vial was replaced by newly-diluted compounds every day. The 4-VA treatment assay was performed by the same method.

To determine the time-dependent effect of 4-VA, control females were treated by paraffin oil from PAE 1-6 days. In parallel, paraffin oil was placed by 4-VA at PAE 1-2 days, 3-4 days, 5-6 days, respectively. The ovaries were dissected and the lengths of terminal oocytes were measured as described above. The brains, CC-CA, and fat body of females were dissected and stored immediately in liquid nitrogen for further experiments.

### Electroantennography (EAG) assays

An aliquot of odor was dissolved in paraffin oil (w/v) and loaded with 10 μl on a 5 × 40 mm filter paper strip (Whatman), which was placed inside a Pasteur pipette. This odor was used on subsequent EAG assay. Hexane was tested as negative controls. The antennae of the adult locusts were cut at the bases of the flagella and distal antennal. Segments were cut off 2 mm and then fixed between two electrodes with electrode gel Spectra 360 (Parker, Orange, N. J. UAS). The EAG signals were amplified, monitored, and analyzed with the EAG-Pro software (IDAC4, Syntech, the Netherlands; EAG software v2.6c). A continuous air flow of 30 ml/s was produced by a stimulus controller (Syntech CS-05). Stimulation duration was 1 s and the intervals were 1 min. The blank was applied at the start and end of the stimulation series. The average EAG amplitude was subtracted from that of the blank.

### Single sensilla responses (SSR) assay

SSRs were recorded and analyzed, and stimuli were prepared as previously described (*Li et al., 2016*). The locust was placed in a plastic tube 1 cm in diameter, and its head and antennae were fixed with dental wax. A tungsten wire electrode was electrolytically sharpened by 10% NaNO_2_. The recording electrode was inserted into the bottom of the sensilla through a micromanipulator (Narishige, Japan). The reference electrode was inserted into the eye. Recording electrodes were connected to amplifiers (IDAC4, Syntech, Netherlands). The frequency variation of each pulse at 0.2 s was calculated by using automatic frequency meter software. Signals were recorded for 10s, starting 1 s before stimulation. The preparation is held in a humidified continuous air flow delivered by the Syntech Stimulus controller (CS-55 model, Syntech) at 1.4 L/min. Chemical substances as SSR stimulants included mineral oil as the blank, which was used to dilute the 4- VA at 1, 10, 100, 1000, 10000, 100000, 1000000 ng/μl, respectively. A piece of filter paper (Whatman, UK) was placed in a 15 cm Pasteur glass tube and 10 μl of volatile solution was added to the filter paper. Responses were calculated by counting the number of action potentials one second after stimulation.

### Total RNA extraction, RNA-seq and Quantitative real-time PCR

Total RNA from different tissues were extracted using the TRIzol reagent (Invitrogen™ TRIzol™ Reagent, Cat.15596026) and treated with DNase I following the manufacturer’s instructions.

For RNA-seq, three independent replicates were performed for each sample. The RNA-seq data reported here have been deposited in the Genome Sequence Archive (Genomics, Proteomics & Bioinformatics 2017) in National Genomics Data Center (Nucleic Acids Res 2020), Beijing Institute of Genomics (China National Center for Bioinformation), Chinese Academy of Sciences, under accession number CRA003038 that are publicly accessible at https://bigd.big.ac.cn/gsa. RNA integrity. cDNA libraries were prepared according to Illumina’s protocols. Raw data were filtered, corrected, and mapped to locust genome sequence using TopHat2 software. The number of total reads was normalized by multiple normalization factors. Transcript levels were calculated using the reads per kb million mapped reads criteria. The differences between the test and control groups were represented by *P* values. DEGs were detected by using edgeR package with significance levels at *P* < 0.05. Principal component analysis (PCA) was accomplished using the princomp and pca functions. Enrichment analysis of the Gene Ontology (GO) was carried out based on an algorithm presented by GOstat.

For qPCR, cDNA was reverse transcribed with 2 μg of total RNA using M-MLV Reverse Transcriptase (Promega, Madison, USA). The relative mRNA levels of targeting genes were quantified by Real Master Mix Kit (Tiangen) with LightCycler 480 instrument (Roche). Melting curve analysis was performed to confirm the specificity of amplification. The primers used for qPCR were presented in Supplementary file 2.

### Protein preparation and Western blot analysis

Ovaries and fat body of 4-VA-treated and control females were collected and homogenized in TRIzol reagent (5 individuals/sample, 6 biological repeats/treatment). Total protein was extracted following manufacturers’ instructions. Total protein (100 μg) were separated by gel electrophoresis and then transferred onto polyvinylidene difluoride membranes (Millipore). Non-specific binding sites on the membranes were blocked with 5% bovine serum albumin. The blots were incubated with the primary antibodies (rabbit anti-Vg serum, 1:500, Beijing Protein Innovation Co., Ltd., BPI) in TBST overnight at 4 °C. After incubation, the membranes were washed, incubated with anti-rabbit IgG secondary antibody (1:5000) (EASYBIO, China) for 1 h at room temperature, and then washed again. Protein bands were detected by chemiluminescence (ECL kit, CoWin). The antibodies were stripped from the blots, re-blocked, and then probed with an anti-GAPDH antibody (1:5000)(*Wang et al., 2013*). Protein bands were detected by chemiluminescence (ECL kit, Thermo Scientific). The intensities of the Western blot signals were quantified using densitometry.

### JH titer measurement

Twenty microliters of hemolymph were added to a 1.5-mL tube with 100 μl of 70% methanol and thoroughly mixed. Then, 200 μl of hexane was added to the solution and thoroughly mixed again. The mixture was centrifuged at 5,000 rcf for 10 min at 4 °C. Then, 150 μl of supernatant was placed into a new tube, and the JH precipitate was dried by nitrogen. The JH precipitate was dissolved in 50% methanol, mixed by vortexing, and centrifuged at 13,000 rpm for 10 min at 4 °C. JHIII in the supernatant was detected using the rapid resolution liquid chromatography system (ACQUITY UPLC I-Class, Waters, USA) An ACQUITY UPLC™ BEH C18 column (50× 2.1 mm, 1.7 μm) was used for LC separation. The autosampler was set at 10 °C, using gradient elution with 0.1% formic acid methanol as solvent A and 0.1% formic acid water as solvent B. The flow rate was set at 0.2 ml/min. Mass spectrometry detection was performed on an AB SCIEX Triple Quad™ 4500 (Applied Biosystems, Foster City, CA, USA) with an electrospray ionization source (Turbo Ionspray). The detection was performed in positive electrospray ionization mode. The [M + H] of the analyte was selected as the precursor ion. The quantitation mode was Multiple Reaction Monitoring (MRM) mode using the mass transitions (precursor ions/product ions). The MRM (m/z) of JHIII was 267.2/235.2. Data acquisition and processing were performed using AB SCIEX Analyst 1.6 Software (Applied Biosystems).

### JH rescue experiment

The active JH analog, s-(+)-methoprene (Santa Cruz Biotech) was topically applied to the pronotum of locusts (150 µg per locust) from PAE 3 days to PAE 4 days according to previously published work (*Song et al., 2013, Wu et al., 2016*), and acetone was used as the control. Meanwhile, the treated females were stimulated with 4-VA. The treated females were dissected at PAE 6 day, and the lengths of terminal oocytes were measured as previously described. The ovaries and fat bodies of females at PAE 4 days were dissected and stored immediately in liquid nitrogen for further western blot experiments.

### Statistical analyses

For the measurement of oviposition time and sexual maturation, individuals were randomly allocated into experimental group and control group, and no restricted randomization was applied. The data that do not meet normal distribution was excluded for the analysis of sexual maturity, mRNA levels, protein levels, as well as JH titer measurement. The distribution of the first oviposition time and the consistency of sexual maturation (represented by the length of terminal oocytes) were analyzed using Levene’s test according to previous studies (*Rohner et al., 2013, He et al., 2016*). The mean value of the first oviposition time between two groups was analyzed using Student’s t-test. One-way ANOVA followed by Tukey’s post-hoc test was used for multi-group comparisons. All data were statistically analyzed using GraphPad Prism 5 software and SPSS 17 software. All experiments were performed with at least three independent biological replicates.

## Acknowledgments

This study was supported by the National Natural Science Foundation of China (Grant NO. 31930012, 31920103004, 31772531, and 32070497) and grants from Chinese Academy of Sciences (nos. 152111KYSB20180036) and Youth Innovation Promotion Association CAS (No. 2021079).

## Supplementary figures and files

**Figure 1-figure supplement 1.**
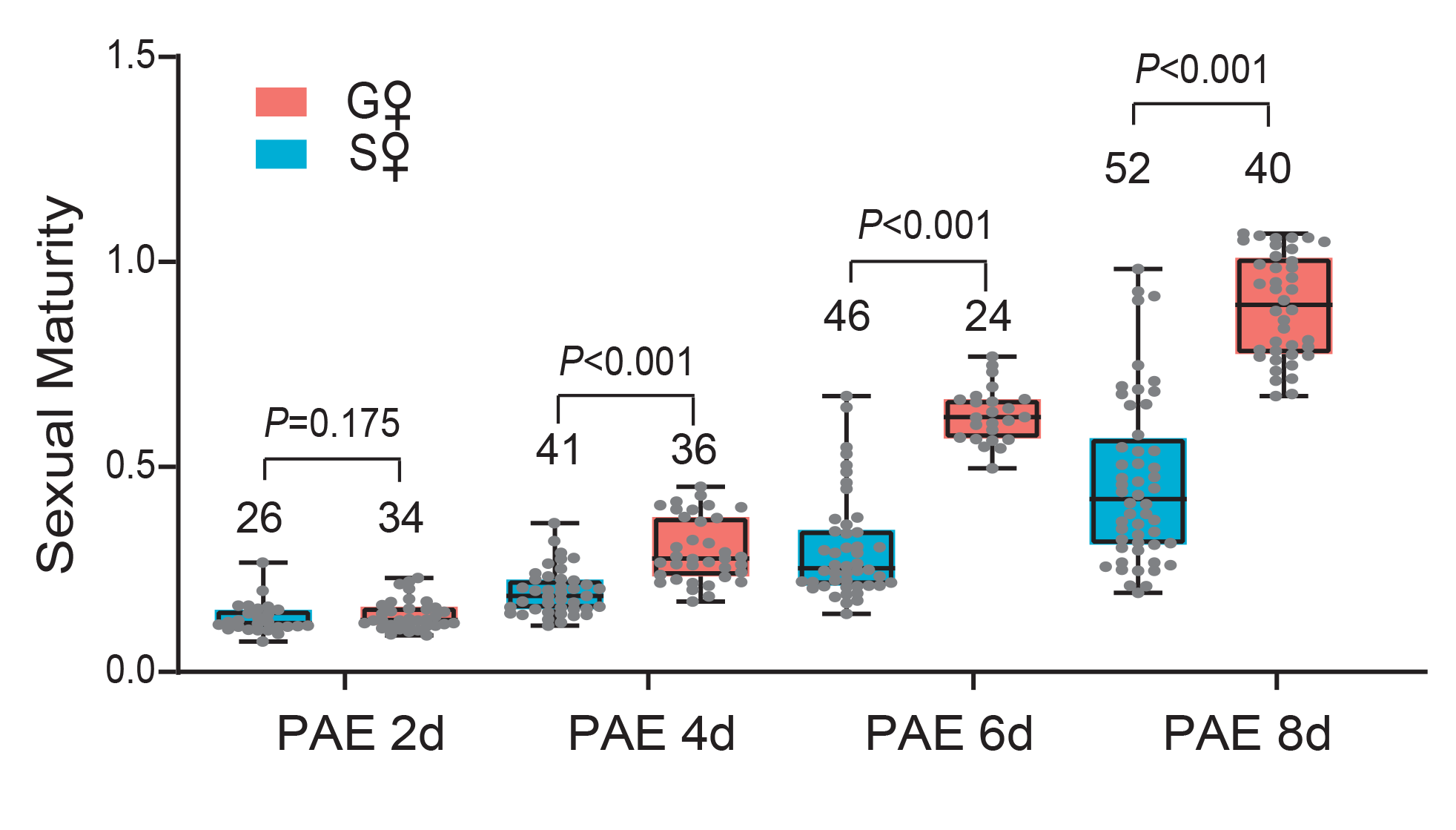
Comparison of sexual maturation rate of gregarious and solitarious female. The maturity of terminal oocytes in gregarious and solitarious females were measured from PAE 2 to PAE 8 days, Data were analyzed using Student’s t-test. The number of biological replicates and *P* values were shown in the figures.

**Figure 1-figure supplement 2.**
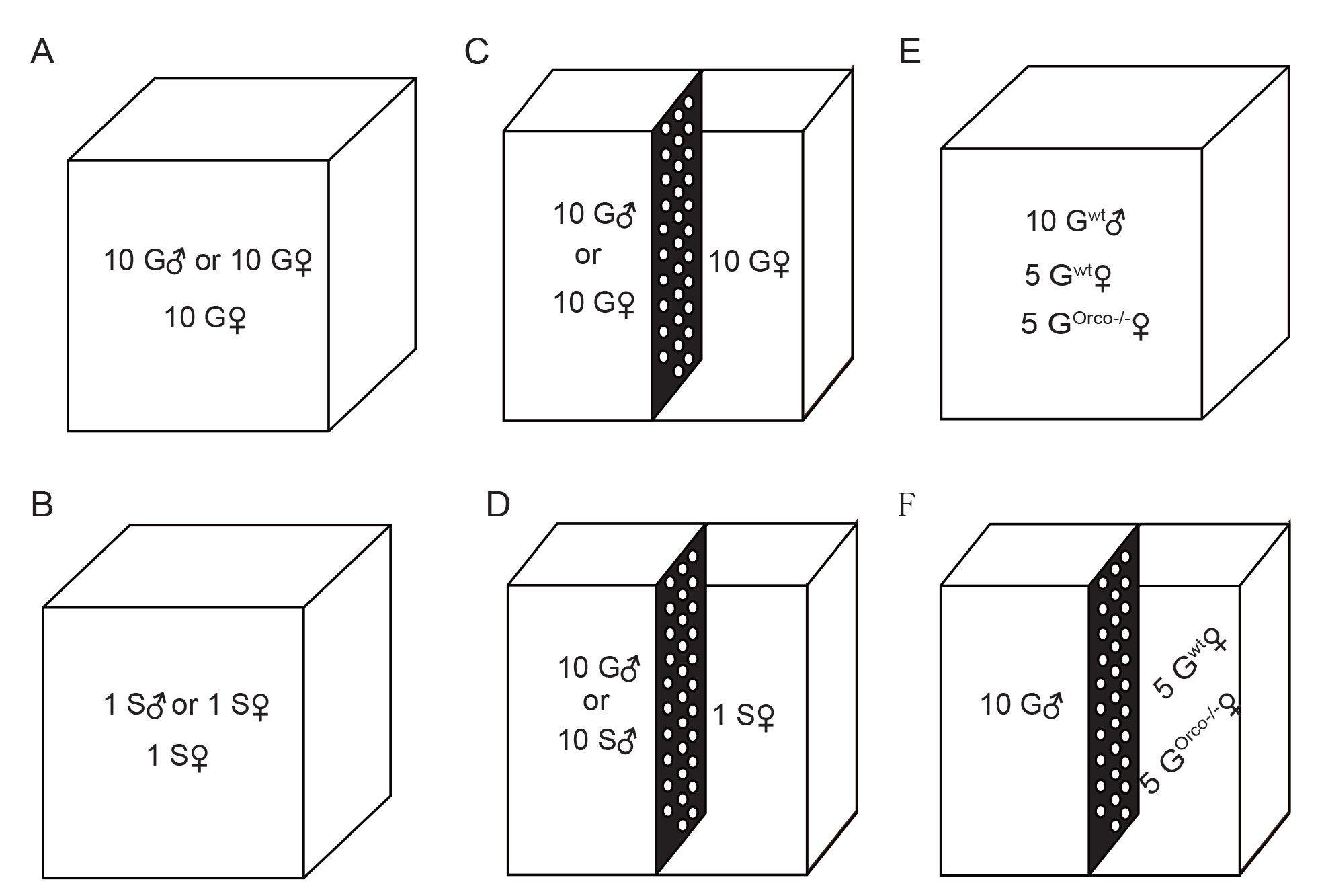
Schematic diagram of the stimulation experiments for females of the migratory locust. (A) Ten gregarious females were reared with ten gregarious males or ten gregarious females after emergence, respectively. (B) One solitarious female was reared with one solitarious male or one solitarious female after emergence, respectively. (C) Ten gregarious females were separately reared with ten gregarious males or ten gregarious females by a breathable partition after emergence. (D) One solitarious female was separately reared with ten gregarious males or ten solitarious males by a breathable partition after emergence. (E) Five Or mutant females (Orco^-/-^ or Or35^-/-^) were reared with five gregarious WT females and ten gregarious WT males from PAE 1-6 days. (F) Five Or mutant females (Orco^-/-^ or Or35^-/-^) and five gregarious WT females were separately reared with ten gregarious WT males by a breathable partition from PAE 1-6 days.

**Figure 2-figure supplement 1.**
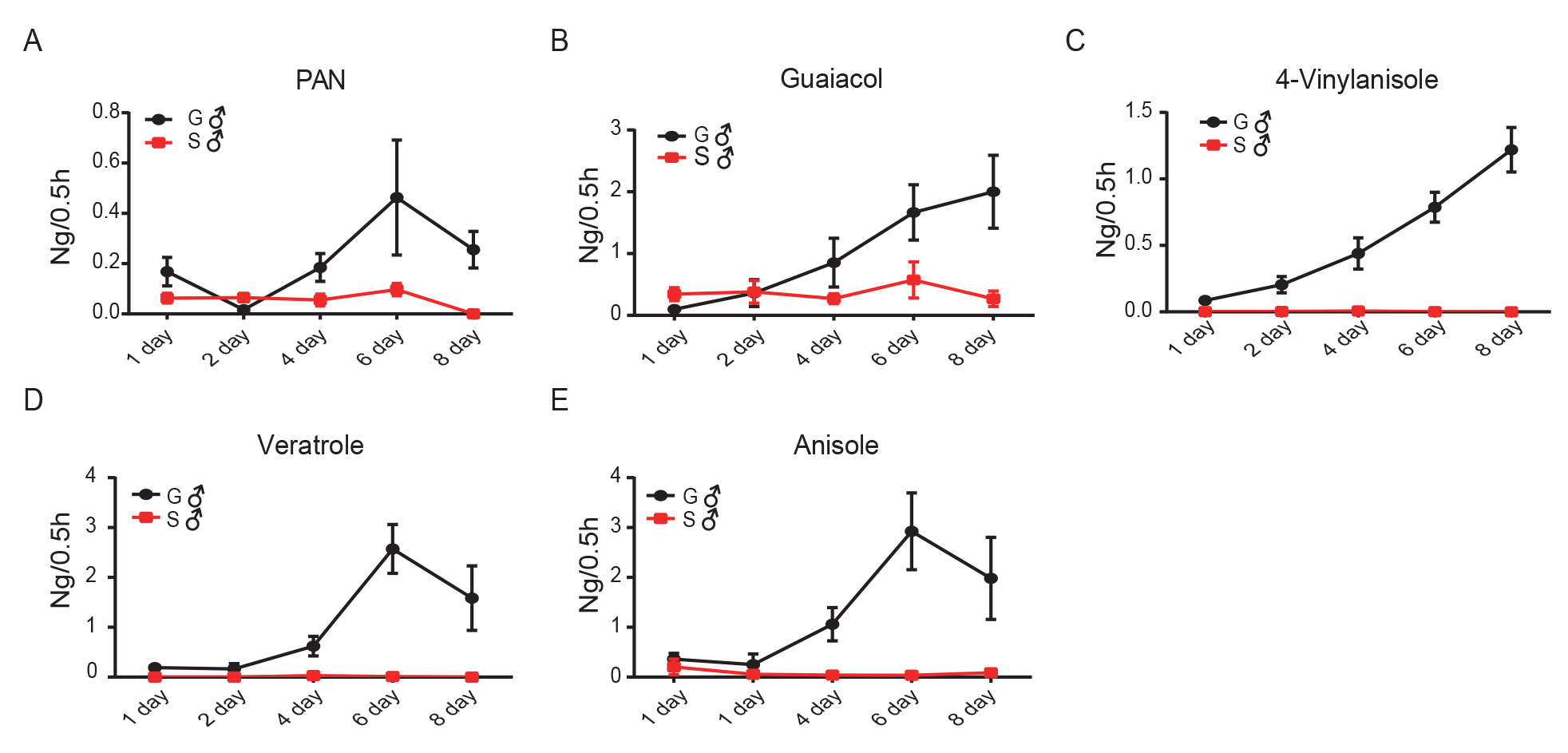
Releasing dynamics of (A) PAN, (B) Guaiacol, (C) 4-VA, (D) Veratrole, (E) Anisole in gregarious and solitarious male adults from PAE 1-8 days. Gregarious male adults release much more PAN, guaicol, 4-vinlanisole, vertrole, and anisole than solitarious male adults after PAE 4 days. Data are shown as means ± SEM (n = 5-8).

**Figure 2-figure supplement 2.**
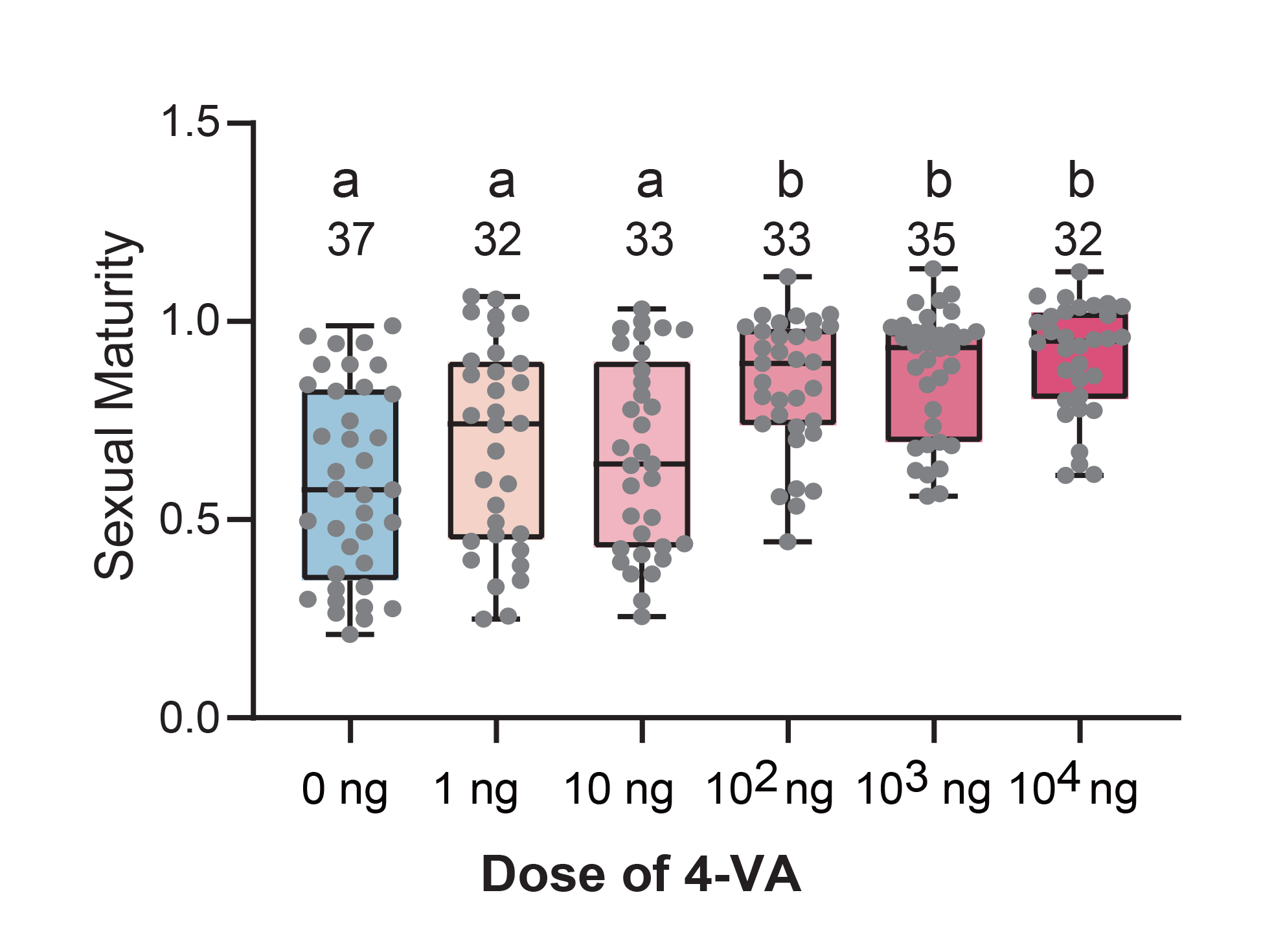
Dose-dependent effects of 4-VA on female maturation rate. Different letters represent significant differences between the two groups (one-way ANOVA, *P* < 0.05). The number of biological replicates were shown in the figures.

**Figure 2-figure supplement 3.**
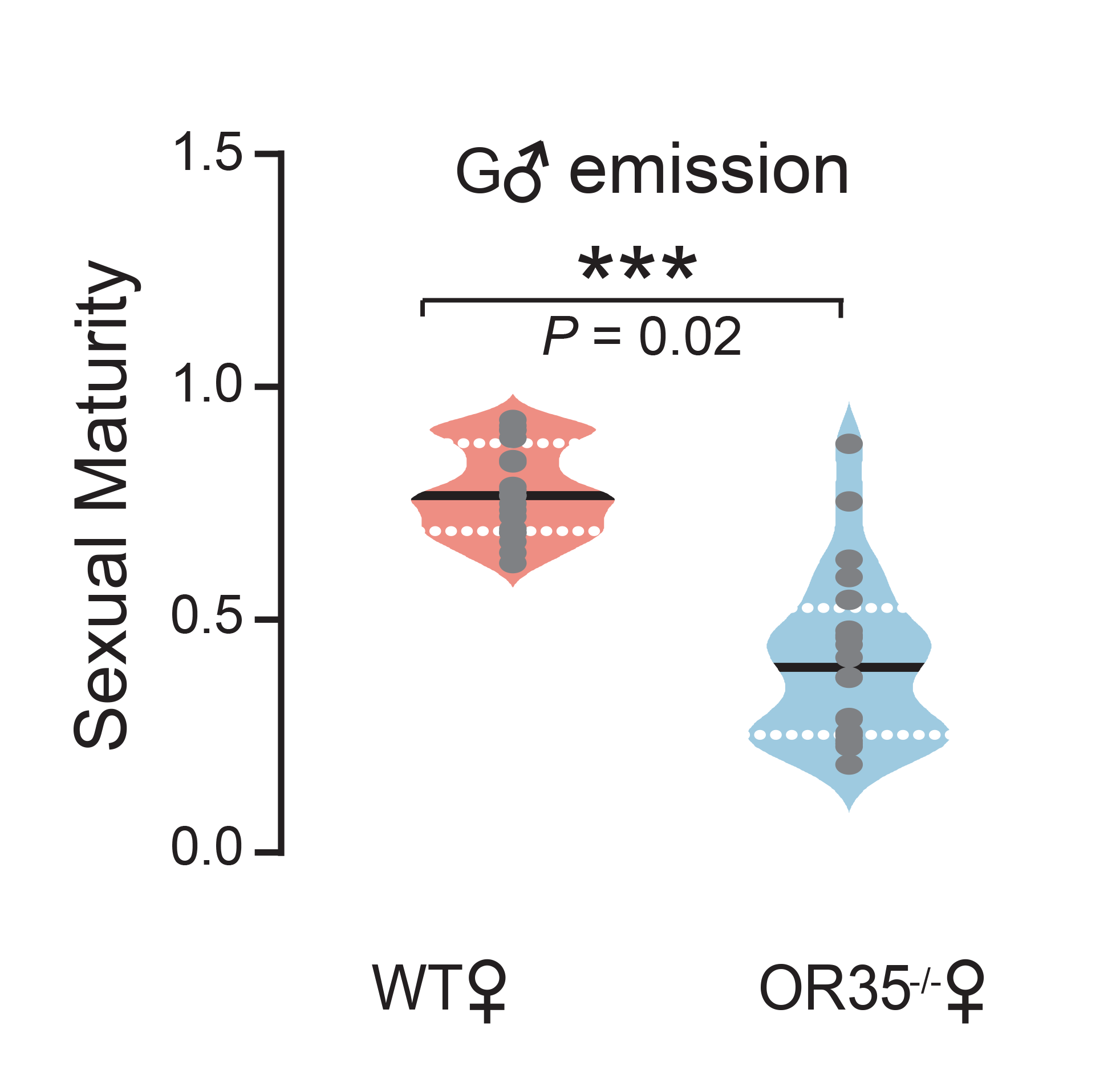
The sextual maturity of WT females and Or35^-/-^ females after stimulation by volatiles released from gregarious males. (n = 20, Levene’s test, *P* = 0.02; Student’s t-test, ***, *P* < 0.001). Lines in droplet diagram indicate median quartile.

**Figure 3-figure supplement 1.**
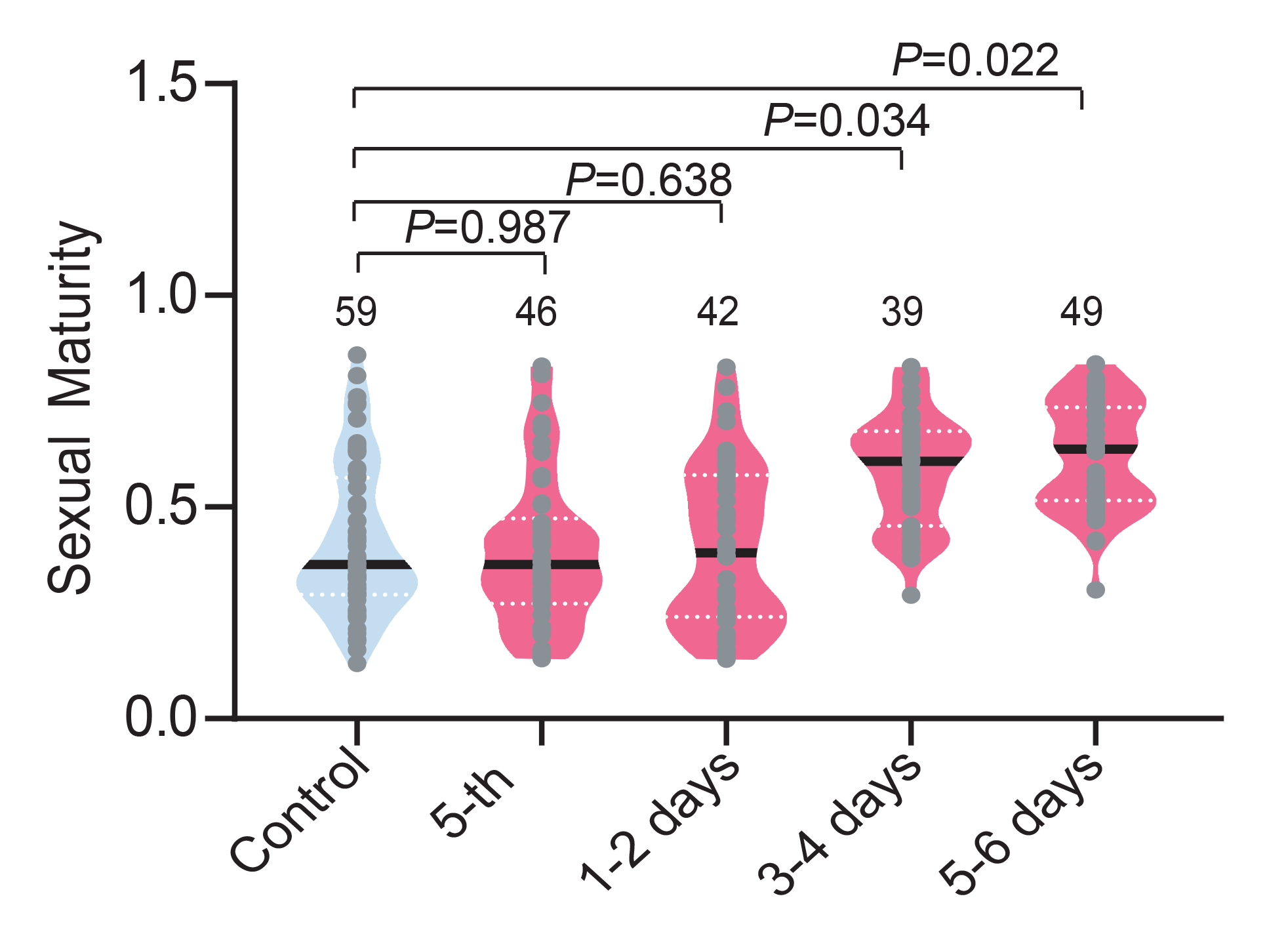
Effects of gregarious males with different ages on maturation synchrony of female aged at PAE3-4 days. Gregarious females aged at PAE 3 days were reared together with the fifth-instar gregarious males, or male adults aged at PAE 1 day, PAE 3 day, PAE 5 day for two days, respectively. Data were analyzed by Levene’s test. The number of biological replicates were shown in the figures. Lines in droplet diagram indicate median quartile.

**Figure 3-figure supplement 2.**
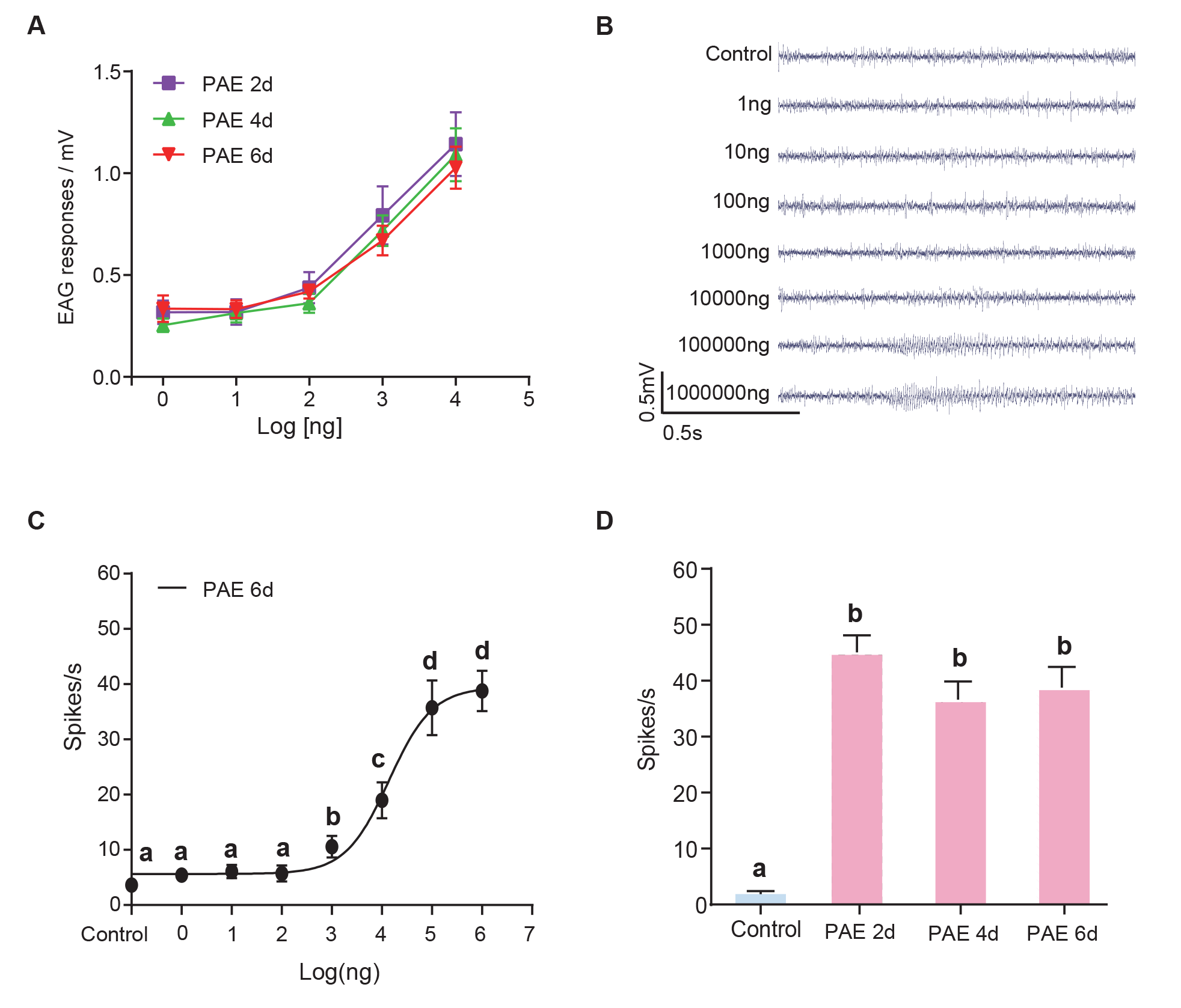
Peripheral electrophysiological responses of female locusts to 4-VA. (A) Dosage effects on EAG responses of females to 4-VA at different developmental stages. Data are shown as means ± SEM. (B) and (C) Dosage effects on single sensilla responses (SSR) of basiconic sensillum in females to 4-VA. (D) The SSR of basiconic sensillum in females to 4-VA aged at different developmental stages. Different letters represent significant differences between the two groups (one-way ANOVA, *P* < 0.05).

**Figure 3-figure supplement 3.**
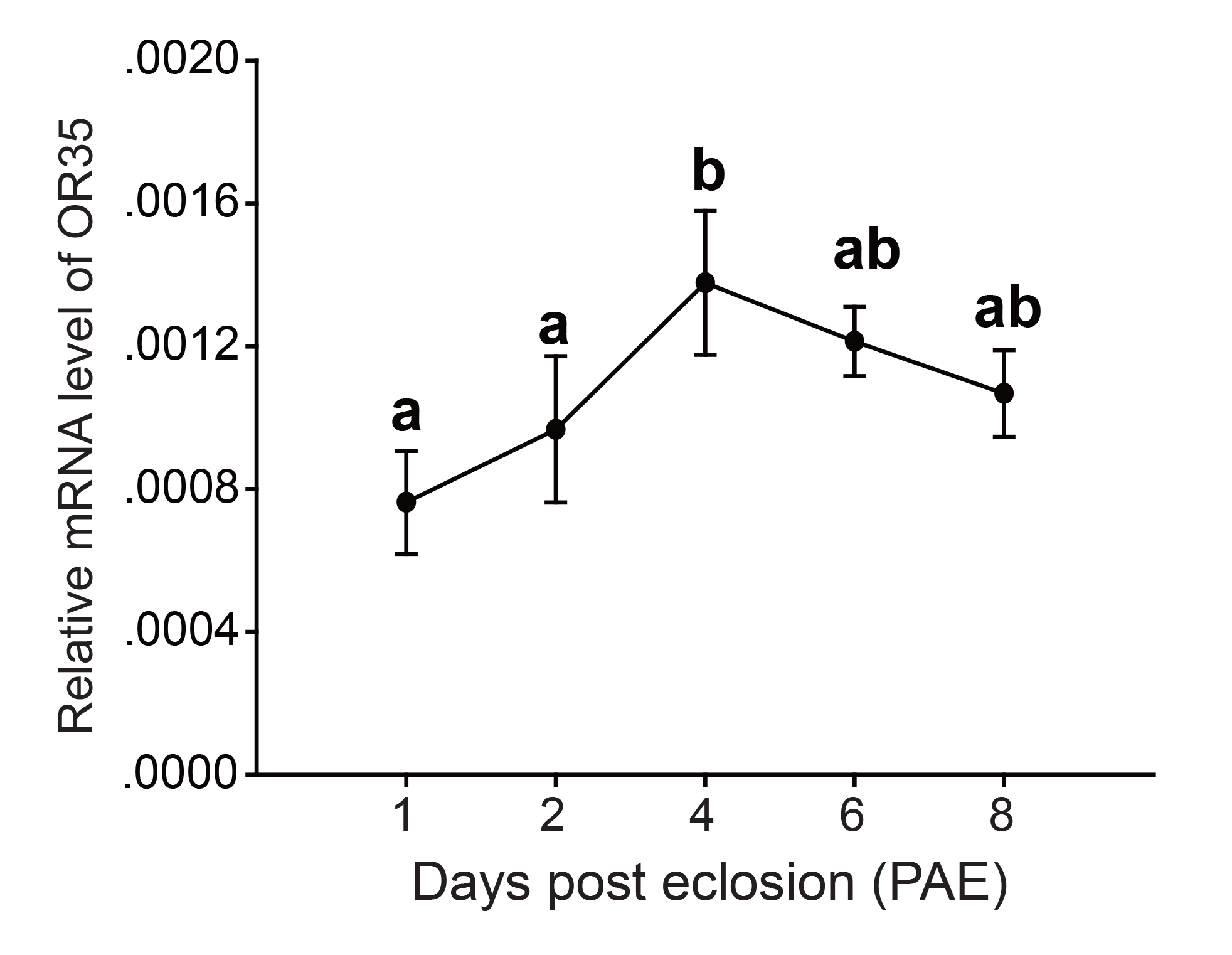
The mRNA levels of *LmigOr35* during PAE 1-8 days. Different letters represent significant differences (one-way ANOVA, *P* < 0.05). Points labeled with different letters indicate a significant difference between these groups. Data are shown as means ± SEM.

**Figure 3-figure supplement 4.**
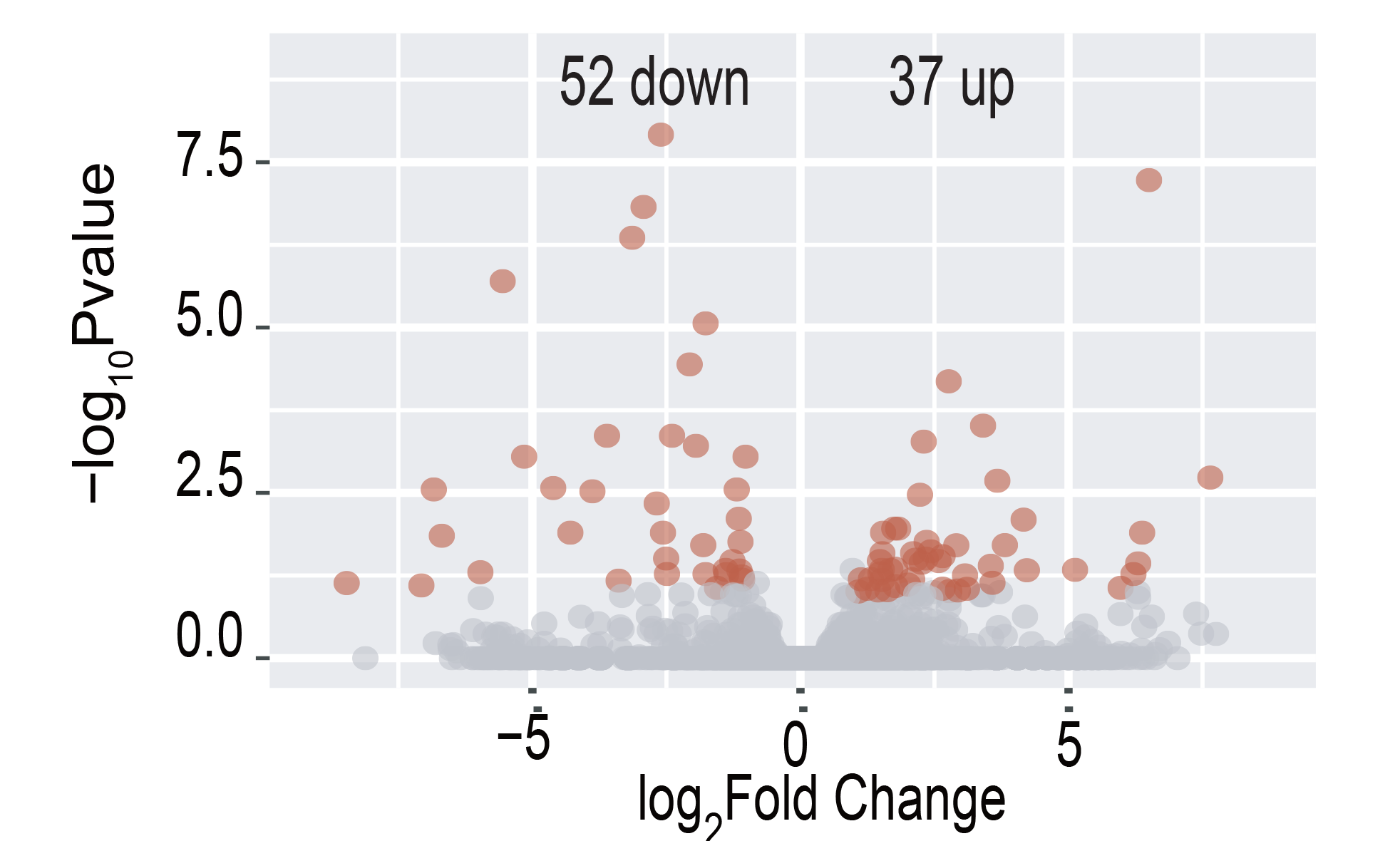
Volcano plot of RNA-seq data in the brain of females treated by 4-VA at PAE 3-4 days. There were 52 down-regulated and 37 up-regulated genes in the brain of female locusts upon 4-VA treatment, respectively.

**Figure 3-figure supplement 5.**
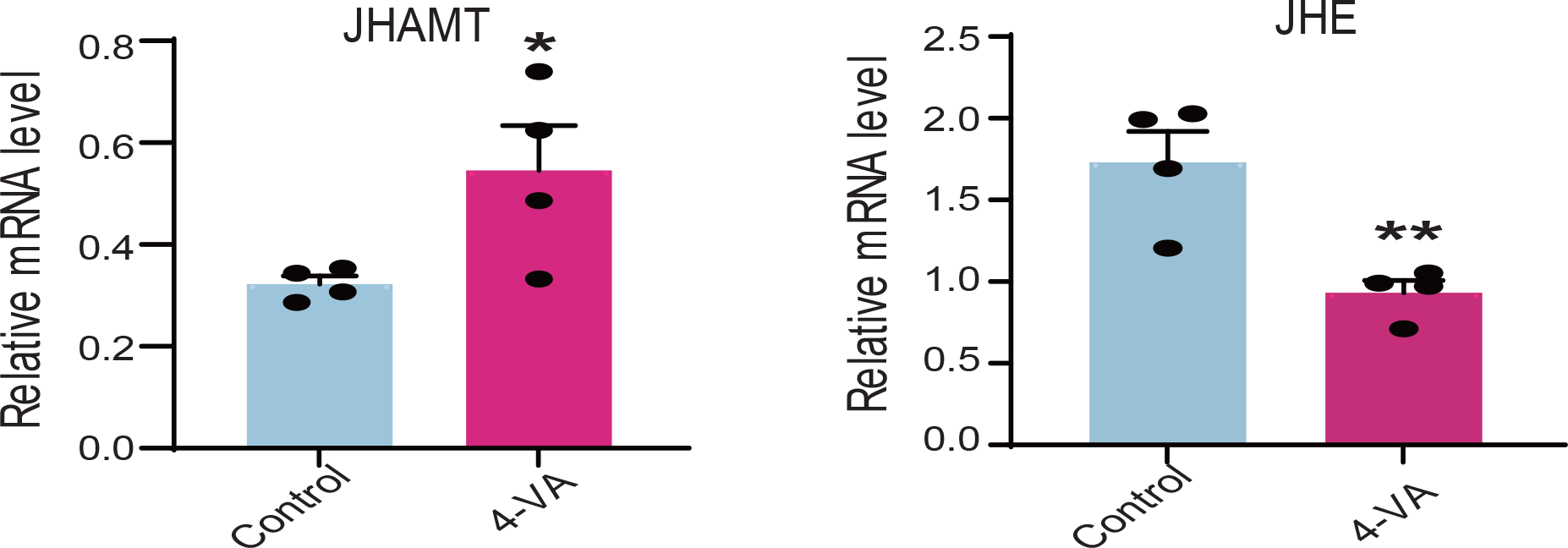
The mRNA level of JHAMT and JHE upon 4-VA treatment. Student’s t-test, n = 4, \**P* < 0.05, \*\**P* < 0.01. Data are shown as means ± SEM.

**Figure 4-figure supplement 1.**
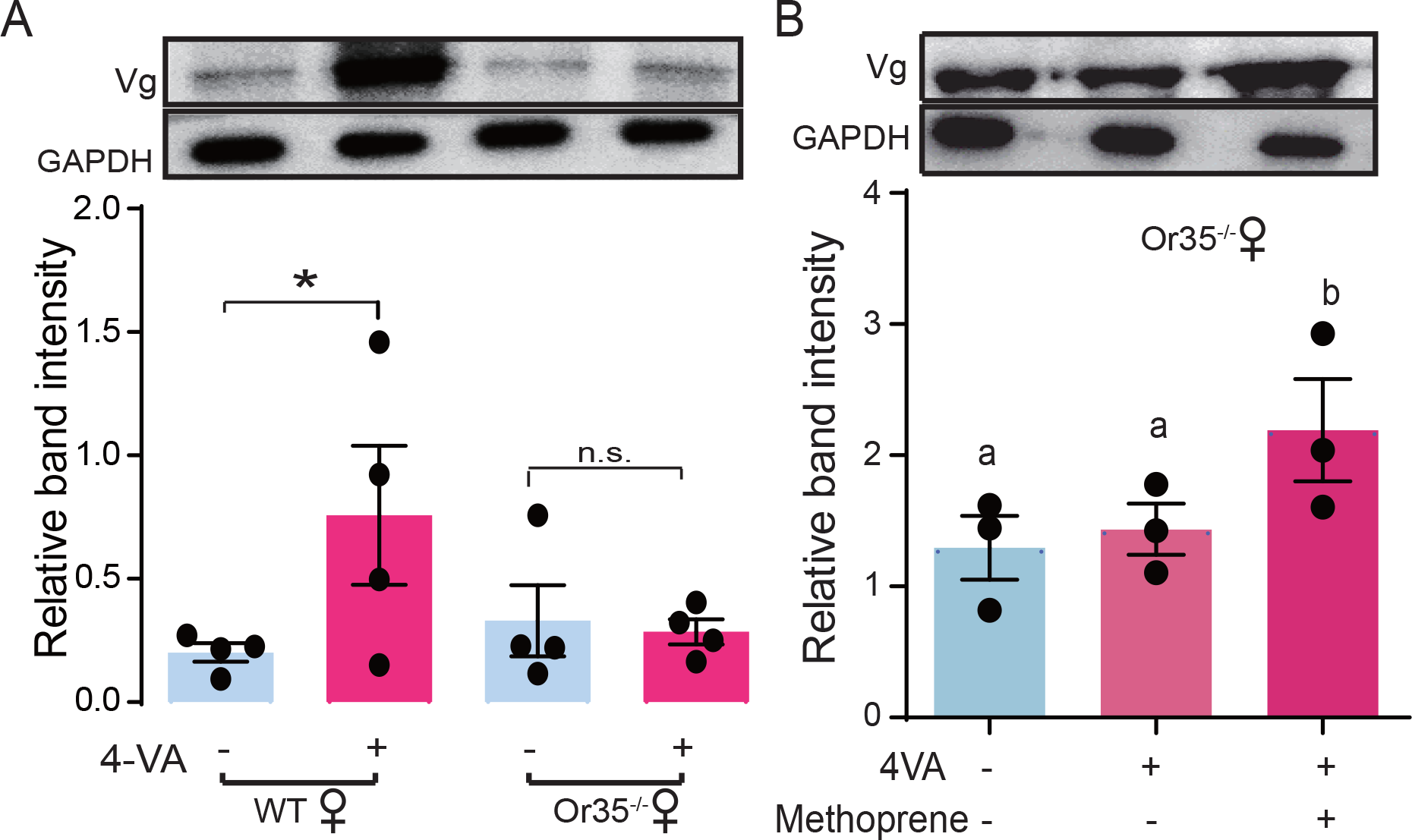
Validation the role of Or35 in 4-VA-enhanced Vg expression in the fat body of female locusts. (A) The protein levels of Vg in the fat body of WT females and Or35^-/-^ females with or without 4-VA treatment. Student’s t-test: n = 4, \**P* < 0.05, n.s. indicates not significant. Data are shown as means ± SEM. (B) The protein levels of Vg in the fat body of WT and Or35^-/-^ females with 4-VA exposure and JHA treatment. Different letters represent significant differences (n = 4, one-way ANOVA, *P* < 0.05). Columns labeled with different letters indicate a significant difference between these groups. Data are shown as means ± SEM.

## Supplementary files

Supplementary file 1. List of genes related to JH synthesis and degradation in the CC-CA of female adults exposure to 4-VA at PAE 3-4 days.

Supplementary file 2. Primers used in qPCR analysis.

## Source data

Figure 1-source data. Raw data for first oviposition time and sexual maturity of gregarious, solitarious, and Orco-/- female adults.

Figure 2-source data. Raw data for volatile contents in male adults and sexual maturity of females.

Figure 3-source data. Raw data for sexual maturity, JH titer, gene expression, and protein level in 4-VA-treated females.

Figure 4-source data. Raw data for JH titer, gene expression, protein level, and sexual maturity in WT and Or35-/- females.

## Supplementary source data

Figure 1-figure supplement 1-source data. Raw data for maturation rate of females between gregarious and solitarious phases.

Figure 2-figure supplement 1-source data. Raw data for volatile contents during male adult development.

Figure 2-figure supplement 2-source data. Raw data for maturation rate of females treated by 4- VA.

Figure 2-figure supplement 3-source data. Raw data for sextual maturity of WT females and Or35-/- females stimulated by volatiles released from gregarious males.

Figure 3-figure supplement 1-source data. Raw data for sextual maturity of females reared with gregarious males with different ages.

Figure 3-figure supplement 2-source data. Raw data for electrophysiological responses of female locusts to 4-VA.

Figure 3-figure supplement 3-source data. Raw data for mRNA levels of LmigOr35 during PAE 1-8 days.

Figure 3-figure supplement 5-source data. Raw data for mRNA level of JHAMT and JHE upon 4- VA treatment.

Figure 4-figure supplement 1-source data. Raw data for Vg expression in the fat body of female locusts stimulated by 4-VA

## References

1. Anstey ML, Rogers SM, Ott SR, Burrows M and Simpson SJ, 2009. Serotonin mediates behavioral gregarization underlying swarm formation in desert locusts. Science 323: 4. DOI: 10.1126/science.1165939.

2. Assad YOH, Hassanali A, Torto B, Mahamat H, Bashir NHH and ElBashir S, 1997. Effects of fifth-instar volatiles on sexual maturation of adult desert locust schistocerca gregaria. Journal of Chemical Ecology 23: 1373–1388. DOI: Doi 10.1023/B:Joec.0000006470.30501.73.

3. Bendena WG, Hui JHL, Chin-Sang I and Tobe SS, 2020. Neuropeptide and microrna regulators of juvenile hormone production. Gen Comp Endocrinol 295: 113507. DOI: 10.1016/j.ygcen.2020.113507.

4. Berger J, 1992. Facilitation of reproductive synchrony by gestation adjustment in gregarious mammals: A new hypothesis. Ecology 73: 323–329.

5. Buck J and Buck E, 1968. Mechanism of rhythmic synchronous flashing of fireflies. Science 159: 1319–1327. DOI: 10.1126/science.159.3821.1319.

6. Cahill LP, Buckmaster JM, Cumming IA, Parr RA and Williams AH, 1974. The effect of the presence of a ram on the time of ovulation in ewes. J Reprod Fertil 40: 475–477.

7. Chen Q, He J, Ma C, Yu D and Kang L, 2015. *Syntaxin 1a* modulates the sexual maturity rate and progeny egg size related to phase changes in locusts. Insect Biochemistry And Molecular Biology 56: 1–8. DOI: 10.1016/j.ibmb.2014.11.001.

8. Dey S, Chamero P, Pru JK, Chien M-S, Ibarra-Soria X, Spencer KR, Logan DW, Matsunami H, Peluso JJ and Stowers L, 2015. Cyclic regulation of sensory perception by a female hormone alters behavior. Cell 161: 1334–1344.

9. Drickamer LC, 1977. Delay of sexual maturation in female house mice by exposure to grouped females or urine from grouped females. Journal Reprod Fertil 51: 5. DOI: 10.1530/jrf.0.0510077.

10. Engel KC, Stokl J, Schweizer R, Vogel H, Ayasse M, Ruther J and Steiger S, 2016. A hormone-related female anti-aphrodisiac signals temporary infertility and causes sexual abstinence to synchronize parental care. Nat Commun 7: 11035. DOI: 10.1038/ncomms11035.

11. French JA and Stribley JA, 1985. Patterns of urinary oestrogen excretion in female golden lion tamarins (leontopithecus rosalia). Journal Reprod Fertil 75: 10. DOI: 10.1530/jrf.0.0750537.

12. Gadenne C, Barrozo RB and Anton S, 2016. Plasticity in insect olfaction: To smell or not to smell? Annu Rev Entomol 61: 317–333. DOI: 10.1146/annurev-ento-010715-023523.

13. Gattermann R, Ulbrich K and Weinandy R, 2002. Asynchrony in the estrous cycles of golden hamsters (mesocricetus auratus). Horm Behav 42: 70–77. DOI: 10.1006/hbeh.2002.1800.

14. Guo F and Xia B, 1964. The male locust secretes substances that promote the maturation of the female locust’s ovaries. Chinese Science Bulletin 01: 3.

15. Guo W, Wang X, Ma Z, Xue L, Han J, Yu D and Kang L, 2011. Csp and takeout genes modulate the switch between attraction and repulsion during behavioral phase change in the migratory locust. PLoS Genet 7: e1001291. DOI: 10.1371/journal.pgen.1001291.

16. Guo W, Wu Z, Song J, Jiang F, Wang Z, Deng S, Walker VK and Zhou S, 2014. Juvenile hormone-receptor complex acts on mcm4 and mcm7 to promote polyploidy and vitellogenesis in the migratory locust. PLoS Genet 10: e1004702. DOI: 10.1371/journal.pgen.1004702.

17. Guo X, Yu Q, Chen D, Wei J, Yang P, Yu J, Wang X and Kang L, 2020. 4-vinylanisole is an aggregation pheromone in locusts. Nature DOI: 10.1038/s41586-020-2610-4.

18. Harrison PL, Babcock RC, Bull GD, Oliver JK, Wallace CC and Willis BL, 1984. Mass spawning in tropical reef corals. Science 223: 1186–1189. DOI: 10.1126/science.223.4641.1186.

19. He J, Chen Q, Wei Y, Jiang F, Yang M, Hao S, Guo X, Chen D and Kang L, 2016. Microrna-276 promotes egg-hatching synchrony by up-regulating brm in locusts. Proc Natl Acad Sci U S A 113: 584–589. DOI: 10.1073/pnas.1521098113.

20. Ims RA, 1990. The ecology and evolution of reproductive synchrony. Trends in ecology & evolution 5: 135–140.

21. Ims RA and Steen H, 1990. Geographical synchrony in microtine population cycles: A theoretical evaluation of the role of nomadic avian predators. Oikos 57: 7. DOI: 10.2307/3565968.

22. Janzen DH, 1971. Seed predation by animals. Annual Review of Ecology and Systematics 2: 28. DOI: 10.1146/annurev.es.02.110171.002341.

23. Jindra M, Palli SR and Riddiford LM, 2013. The juvenile hormone signaling pathway in insect development. Annu Rev Entomol 58: 181–204. DOI: 10.1146/annurev-ento-120811-153700.

24. Jovani R and Grimm V, 2008. Breeding synchrony in colonial birds: From local stress to global harmony. Proc Biol Sci 275: 1557–1563. DOI: 10.1098/rspb.2008.0125.

25. Kelly D and Sork VL, 2002. Mast seeding in perennial plants: Why, how, where? Annual Review of Ecology and Systematics 33: 427–447. DOI: 10.1146/annurev.ecolsys.33.020602.095433.

26. Kobayashi M, Sorensen PW and Stacey NE, 2002. Hormonal and pheromonal control of spawning behavior in the goldfish. Fish Physiology and Biochemistry 26: 71–84. DOI: Doi 10.1023/A:1023375931734.

27. Korb J, 2015. Juvenile hormone: A central regulator of termite caste polyphenism. Advances in Insect Physiology 48: 131–161.

28. Li Y, Zhang J, Chen D, Yang P, Jiang F, Wang X and Kang L, 2016. Crispr/cas9 in locusts: Successful establishment of an olfactory deficiency line by targeting the mutagenesis of an odorant receptor co-receptor (orco). Insect Biochem Mol Biol 79: 27–35. DOI: 10.1016/j.ibmb.2016.10.003.

29. Loher W, 1960. The chemical acceleration of the maturation process and its hormonal control in the male of the desert locust.

30. Loher W, 1961. The chemical acceleration of the maturation process and its hormonal control in the male of the desert locust. Proceedings of the Royal Society B: Biological Sciences 153: 380–397. DOI: 10.1098/rspb.1961.0008.

31. Luo M, Li D, Wang Z, Guo W, Kang L and Zhou S, 2017. Juvenile hormone differentially regulates two grp78 genes encoding protein chaperones required for insect fat body cell homeostasis and vitellogenesis. J Biol Chem 292: 8823–8834. DOI: 10.1074/jbc.M117.780957.

32. Mahamat H, Hassanali A and Odongo H, 2000. The role of different components of the pheromone emission of mature males of the desert locust, schistocerca gregaria (forskål)(orthoptera: Acrididae) in accelerating maturation of immature adults. International Journal of Tropical Insect Science 20: 1–5.

33. Mahamat H, Hassanali A, Odongo H, Torto B and EI-Bashir E, 1993. Studies on the maturation- accelerating pheromone of the desert locust schistocerca gregaria (orthoptera: Acrididae). Chemoecology 4: 159–164.

34. McClintock MK, 1978. Estrous synchrony and its mediation by airborne chemical communication (rattus norvegicus). Hormones and Behavior 10: 13. DOI: 10.1016/0018-506X(78)90071-5.

35. Nishide Y and Tanaka S, 2016. Desert locust, schistocerca gregaria, eggs hatch in synchrony in a mass but not when separated. Behavioral Ecology and Sociobiology 70: 1507–1515. DOI: 10.1007/s00265-016-2159-2.

36. Noguera JC and Velando A, 2019. Bird embryos perceive vibratory cues of predation risk from clutch mates. Nature Ecology & Evolution DOI: 10.1038/s41559-019-0929-8.

37. Norris MJ, 1952. Reproduction in the desert locust (schistocerca gregaria forsk.) in relation to density and phase. Anti-Locust Bulletin 49 pp.-49 pp.

38. Norris MJ, 1954. Sexual maturation in the desert locust (schistocerca gregaria forsk. ) with special reference to the effects of grouping. Anti-Locust Bull. 28: 27.

39. Norris MJ, ; Richards, O., 1964. accelerating and inhibiting effects of crowding on sexual maturation in two species of locusts.Pdf. Nature 203: 784–785.

40. Norris MJ and Richards O, 1964. Accelerating and inhibiting effects of crowding on sexual maturation in two species of locusts. Nature 203: 784–785.

41. Nouzova M, Rivera-Perez C and Noriega FG, 2018. Omics approaches to study juvenile hormone synthesis. Curr Opin Insect Sci 29: 49–55. DOI: 10.1016/j.cois.2018.05.013.

42. Odhiambo TR, 1966. Growth and the hormonal control of sexual maturation in the male desert locust schistocerca gregaria. Trans. R. ent. SOC. Lond 118: 393–412.

43. Pener MP and Simpson SJ (2009). Locust phase polyphenism: An update. <u>Advances in insect</u> <u>physiology</u>. S. J. Simpson and M. P. Pener, Academic Press. 36: 1–272.

44. Rekwot P, Ogwu D, Oyedipe E and Sekoni V, 2001. The role of pheromones and biostimulation in animal reproduction. Animal reproduction science 65: 157–170.

45. Robinson GE and Vargo EL, 1997. Juvenile hormone in adult eusocial hymenoptera: Gonadotropin and behavioral pacemaker. Arch Insect Biochem Physiol 35: 559–583. DOI: 10.1002/(SICI)1520-6327(1997)35:4<559::AID-ARCH13>3.0.CO;2-9.

46. Rohner N, Jarosz DF, Kowalko JE, Yoshizawa M, Jeffery WR, Borowsky RL, Lindquist S and Tabin CJ, 2013. Cryptic variation in morphological evolution: Hsp90 as a capacitor for loss of eyes in cavefish. Science 342: 1372–1375. DOI: 10.1126/science.1240276.

47. Song J, Wu Z, Wang Z, Deng S and Zhou S, 2014. Kruppel-homolog 1 mediates juvenile hormone action to promote vitellogenesis and oocyte maturation in the migratory locust. Insect Biochem Mol Biol 52: 94–101. DOI: 10.1016/j.ibmb.2014.07.001.

48. Song JS, Guo W, Jiang F, Kang L and Zhou ST, 2013. Argonaute 1 is indispensable for juvenile hormone mediated oogenesis in the migratory locust, locusta migratoria. Insect Biochem Molec 43: 879–887. DOI: 10.1016/j.ibmb.2013.06.004.

49. Torto B, Njagi PG, Hassanali A and Amiani H, 1996. Aggregation pheromone system of nymphal gregarious desert locust,schistocerca gregaria (forskal). J Chem Ecol 22: 2273–2281. DOI: 10.1007/BF02029546.

50. Torto B, Obeng-Ofori D, Njagi PG, Hassanali A and Amiani H, 1994. Aggregation pheromone system of adult gregarious desert locust schistocerca gregaria (forskal). J Chem Ecol 20: 1749–1762. DOI: 10.1007/BF02059896.

51. Uvarov BP (1977). Grasshoppers and locusts: A handbook of general acridology. London, UK: Centre for Overseas Pest Research, Cambridge University Press.

52. Uzsak A and Schal C, 2012. Differential physiological responses of the german cockroach to social interactions during the ovarian cycle. J Exp Biol 215: 3037–3044. DOI: 10.1242/jeb.069997.

53. Vandenbergh JG, 1967. The development of social structure in free-ranging rhesus monkeys. Behaviou 29: 16. DOI: 10.2307/4533189.

54. Wang X and Kang L, 2014. Molecular mechanisms of phase change in locusts. Annu Rev Entomol 59: 225–244. DOI: 10.1146/annurev-ento-011613-162019.

55. Wang YD, Yang PC, Cui F and Kang L, 2013. Altered immunity in crowded locust reduced fungal (metarhizium anisopliae) pathogenesis. Plos Pathog 9: DOI: ARTN e100310210.1371/journal.ppat.1003102.

56. Ward A and Webster M (2016). The evolution of group living. Sociality: The behaviour of group-living animals: 191–216.

57. Wei J, Shao W, Wang X, Ge J, Chen X, Yu D and Kang L, 2017. Composition and emission dynamics of migratory locust volatiles in response to changes in developmental stages and population density. Insect Sci 24: 60–72. DOI: 10.1111/1744-7917.12396.

58. Wu Z, Guo W, Yang L, He Q and Zhou S, 2018. Juvenile hormone promotes locust fat body cell polyploidization and vitellogenesis by activating the transcription of cdk6 and e2f1. Insect Biochem Mol Biol 102: 1–10. DOI: 10.1016/j.ibmb.2018.09.002.

59. Wu ZX, Guo W, Xie YT and Zhou ST, 2016. Juvenile hormone activates the transcription of cell-division-cycle 6 (cdc6) for polyploidy-dependent insect vitellogenesis and oogenesis. J Biol Chem 291: 5418–5427. DOI: 10.1074/jbc.M115.698936.

